# Immunogenomic profiling of circulating T cells in pediatric cancer patients during standard-of-care

**DOI:** 10.1101/2025.01.10.632468

**Authors:** Arash Nabbi, Yiyue Jiang, Osvaldo Espin Garcia, Suluxan Mohanraj, Stephanie Pedersen, Johanna Regala, Lauren Vernau, Jessica Perazzelli, David M. Barrett, Trevor J. Pugh

**Affiliations:** Princess Margaret Cancer Centre, University Health Network, Toronto, Ontario, Canada; Department of Medical Biophysics, University of Toronto, Toronto, Ontario, Canada; Center for Childhood Cancer Research Cell Therapy Program, Children’s Hospital of Philadelphia, Philadelphia, Pennsylvania, USA; Ontario Institute for Cancer Research, Toronto, Ontario, Canada; Kite Pharma, Santa Monica, CA, USA

## Abstract

While pediatric cancer patients receive intensive chemotherapy, its impact on peripheral T cells and subsequently to disease outcomes are not fully characterized. Here, we assessed T-cell dynamics during treatment, identifying associations with outcomes through immune phenotyping and T-cell Receptor (TCR) sequencing in pediatric solid and hematologic malignancies. We show that while levels of immune checkpoint proteins (PD-1, LAG3, and TIM3) at baseline were highest in lymphomas compared to other cancer groups, they increased significantly in response to therapy in all cancers. Levels of Central Memory (CM) T cells increased in leukemias and solid tumors, while naïve T cells and cell-free TCR diversity decreased in lymphomas. By combining immune cell and TCR repertoire features across all timepoints, we proposed the Dynamic Immunogenomic Score (DIS) to measure patient-specific effects of therapy on the peripheral T-cell population. Higher DIS was associated with high-risk cancer types and logistic regression analysis revealed it may predict incidence of relapse in leukemia patients. TCR specificity analysis revealed patient-specific clonal dynamics and differential detection of virally-associated TCRs in cancer patients compared to healthy individuals. Our results highlight the potential of early upfront immunogenomic profiling in identifying high-risk patients that may be predictive in light of emerging cellular immunotherapies.

## Introduction

Pediatric cancer patients exhibit variable immune competency as a result of prolonged exposure to chemotherapy regimens. In addition to well-known immediate effects such as neutropenia and risk of infection^1–3^, chemotherapy impacts the immune reconstitution of T cells to varying degrees^4^. For instance, in pediatric acute lymphoblastic leukemia (ALL), levels of total lymphocytes remain low during and after chemotherapy, with levels of CD4+ T cells recovering at a slower rate compared to their CD8+ counterparts^5^. Similarly, in patients with sarcoma or non-Hodgkin lymphoma, intensive chemotherapy reduces the levels of CD4+ and, to a lesser degree, CD8+ T cells ^6^. In a study of patients with brain tumors, sarcomas, and non-hodgkin’s lymphomas, while CD8+ T cells show early recovery three months post-therapy, CD4+ T cells persist below baseline levels^7^. In a basket cohort of 43 children with various types of solid and hematological malignancies, alterations in peripheral T cells or immunoglobulins persisted in > 80% of cases 9-12 months after therapy completion^8^. Similar results were recently reported, showing that in cancer survivors of solid tumors, levels of CD4+ T cells were lower than those in normal controls even 12-60 months after therapy completion^9^. Immunophenotyping of 31 children with various hematological malignancies showed that levels of memory T cells remained low even 5 years post-chemotherapy ^10^. These findings revealed the immunological alterations induced by chemotherapy in pediatric cancer patients, potentially impacting disease outcomes. Indeed, one study showed that Absolute Lymphocyte Count (ALC) at the end of induction therapy was an independent predictor of disease relapse and overall survival in ALL patients^11^. High absolute lymphocyte and monocyte count ratio post-transplantation in relapsed/refractory (R/R) Hodgkin lymphoma patients showed higher 4-year relapse-free survival ^12^. However, the prognostic value of T-cell dynamics during initial therapy remains uncharacterized.

Understanding the impact of standard therapy on the immune system is of clinical value for pediatric cancer patients, as it may improve patient assignment to different types of immunotherapeutic trials. Early trials investigating the Immune Checkpoint Blockers (ICB) in pediatric cancers showed limited efficacy ^13–15^. However, these patients had received multiple prior chemotherapy regimens, possibly rendering their immune system dysfunctional. Accordingly, the T-cell clonal expansion and shrinkage as a result of chemotherapy that may reveal TCRs reactive to tumor antigens or immunogenic cell death, is currently less investigated. In the context of Chimeric Antigen Receptor (CAR) T-cell therapies, patients’ peripheral T cells are used to manufacture CAR products and prolonged chemotherapy exposure may hinder T-cell expansion and persistence of the final CAR product, which are critical for the favorable outcome ^16–18^. Therefore, early identification of at-risk patients together with ideal blood samples for CAR manufacturing informs precision cellular immunotherapy, particularly as CAR T-cell therapies are now being investigated for high-risk patients with post-consolidation minimal residual disease (AALL1721/Cassiopeia)^19,20^.

Here, we sought to determine dynamics of T cells in pediatric hematological and solid malignancies during first-line therapy using flow cytometry and TCR sequencing. Combining immunophenotyping and TCR sequencing features showed an association with the incidence of relapse in leukemia, highlighting the potential value of immune dynamics as a prognostic biomarker during standard treatment. Our study showcases the value of immunogenomic profiling as a dynamic biomarker that can be used to tailor care according to risk of relapse.

## Results

### Pediatric cancer standard-of-care cohort

To investigate changes in the circulating T-cell repertoire during the course of chemoradiation, we collected serial blood samples from119 children beginning with diagnosis of a primary hematological malignancy or solid tumor. This cohort includes 55 leukemias [Acute Lymphoblastic Leukemia (ALL, n = 23), High Risk ALL (HR ALL, n = 14), Standard Risk ALL (SR ALL, n = 3), Acute Myeloid Leukemia (AML, n = 11), Chronic Myeloid Leukemia (CML, n = 4)], 19 lymphomas [Hodgkin’s Disease (HD, n = 8), Burkitt Lymphoma (BL, n = 4), Diffuse Large B-cell Lymphoma (DLBCL, n = 3), B Lymphoblastic Lymphoma (BLL, n = 2)], 39 solid tumors [Ewing Sarcoma (EWS, n = 6), Osteosarcoma (OS, n = 12), Neuroblastoma (NB, n = 5, one metastatic), Hepatoblastoma (HB, n = 4), Alveolar Rhabdomyosarcoma (ARMS, n = 1), Embryonal Rhabdomyosarcoma (ERMS, n = 5), and Wilms tumor (WT, n = 3)], and 6 T-cell malignancies [T-cell Acute Lymphoblastic Leukemia (T-ALL, n = 4), Anaplastic Large Cell Lymphoma (ALCL, n = 2)] (FigS1).

Among the 119 patients enrolled in our study, 23 (19%) subsequently relapsed (8 with AML, 5 with HR ALL, 4 with OS, and single cases in CML, ARMS, EWS, HD, NB, and T-ALL). Patients with relapsed AML underwent bone marrow transplantation, with two achieving remission. The 5 relapsed HR ALL patients were treated with CAR T-cell therapy (Kymriah or humanized CTL119) resulting in remission in 3 patients and lineage switch to AML in 2 patients. The 4 patients with relapsed OS exhibited pulmonary recurrence with 2 remaining in remission post-resection (FigS1, TableS1).

### Immune cell and T-cell repertoire profiling of serial blood collections

To characterize changes in T-cell functional states and repertoire over the course of treatment, we collected blood draws before each chemotherapy cycle for up to five cycles. From 455 PBMC samples from 118 patients, we used flow cytometry to establish the frequency of five T-cell populations, Naïve (CCR7^+^, CD62L^+^, CD45RO^−^, and CD95^−^), Stem-like Central Memory (SCM, CCR7^+^, CD62L^+^, CD45RO^−^, and CD95^+^), Central Memory (CM, CCR7^+^, CD62L^+^, CD45RO^+^, and CD95^+^), Effector Memory (EM, CCR7^−^, CD62L^−^, CD45RO^+^, and CD95^+^), and Terminal Effector (TE, CCR7^−^, CD62L^−^, CD45RO^−^, and CD95^+^). We complemented our analysis by profiling three immune checkpoint proteins, TIM3, LAG3, and PD1 on the surface of CD3+ cells. To characterize T-cell repertoires in this cohort, we performed the CapTCR-seq hybrid capture method followed by deep sequencing^21^ of 197 PBMC and 266 cell-free DNA (FigS1, TableS1).

### Lymphomas expressed high levels of immune checkpoint proteins and harbored a high percentage of Central Memory T cells

We first sought to determine systematic difference in the expression levels of PD1, TIM3, and LAG3, as the primary target for Immune Checkpoint Blockers (ICB), in circulating T cells across pediatric cancers.

In pre-treatment lymphoma PBMC samples, we found higher percentages of PD1, TIM3, and LAG3 compared to those obtained from pre-treatment leukemias or solid tumors (PD1 medians 28.8, 9.4, and 4.3, two-sided Kolmogorov–Smirnov test, p = 0.002 and 0.0008; TIM3 medians 28.9, 9.4, and 4.3, two-sided Kolmogorov–Smirnov test, p = 0.0008 and 0.001; LAG3 medians, 16.7, 7, and 4.7, two-sided Kolmogorov–Smirnov test, p = 0.006 and 0.02) (Fig1A-C). These included samples from three Burkitt lymphoma patients with the highest percentage of T cells expressing PD1 (PD1%) and LAG3 (LAG3%) (medians, 30.67% and 25%), in line with a previous report^22^.

**Figure 1.**
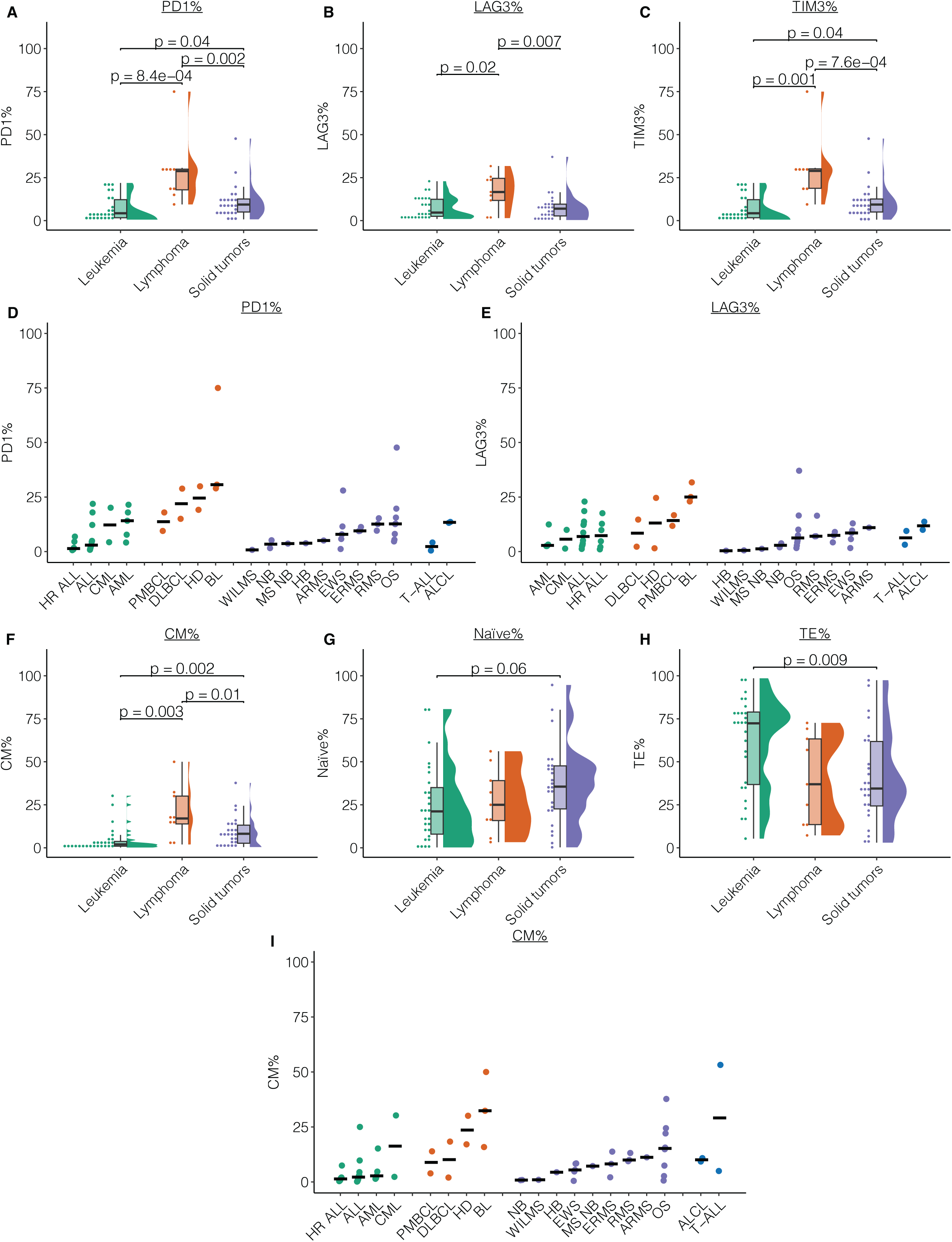
Immunogenomic profiling of peripheral T cells reveals baseline differences among cancer groups. **A-C)** Rain cloud plots showing PD1% (A), LAG3% (B), and TIM3% (C) on the peripheral T cells pre-therapy across three pediatric cancer groups (two-sided Kolmogorov–Smirnov test, N = 59 patients). **D-E)** Breakdown of PD1% (D) and LAG3% (E) (y-axes) by cancer type (x-axes). Cancer types are ordered by their median (black bar) within their cancer groups. **F-H)** Rain cloud plots showing CM% (F), Naïve% (G), and TE% (H) pre-therapy across three pediatric cancer groups (two-sided Kolmogorov–Smirnov test, N = 61 patients). **I)** Breakdown of CM% (y-axis) by cancer type (x-axis). Cancer types are ordered by their median (black bar) within their cancer groups. In boxplots, boxes show median and interquartile range (IQR) and whiskers represent 1.5 × IQR.

Within the pre-treatment leukemia group, peripheral T cells from AML patients had the highest PD1%, while HR ALL had the lowest PD1% in pre-treatment samples (medians, 14.1% and 1.35%, Fig1D). Conversely, AML blood samples had lower LAG3% compared to HR ALL (medians, 2.8% and 7.3%, Fig1E).

In solid tumors, we found lowest levels of PD1% and LAG3% in WT, NB, and HB cases, while OS harbored highest median PD1% and EWS the highest median of LAG3% (medians, 12.6% and 8.5%) (Fig1D-E). Notably, in an OS patient (CHP_390) who relapsed with an isolated pulmonary nodule, we found highest levels of PD1% (47.6%) and LAG3% (37%) in peripheral T cells isolated from pre-treatment samples (Fig1D-E). Collectively, these results reveal overall elevated expression of immune checkpoint proteins in circulating T cells in pre-treatment lymphoma samples, potentially as one of the mechanisms of favorable clinical benefit observed in lymphoma patients treated with ICB, as well as variability across and within pediatric cancer groups.

We next sought to determine whether the differential levels of immune checkpoint proteins were associated with levels of T-cell subsets across pediatric cancers. In lymphomas, CM% was higher compared to leukemias and solid tumors (medians, 17%, 2%, and 8,2%, respectively, two-sided Kolmogorov–Smirnov test, p = 0.003 and p = 0.01, Fig1F). We noted trends of CM% levels across cancer groups generally consistent with those of PD1%, with HR ALL harboring lowest levels across leukemia (median, 0.74%), Burkitt’s lymphoma showing highest levels within lymphomas (median, 24.1%), and OS showing the highest levels within solid tumors (median, 20%) (Fig1I). Notably, patient CHP_390 harbored the lowest levels of CM% within the OS cohort (0.6%), while harboring outlier high levels of SCM% (55.14%, FigS2A). We found otherwise no significant differences among pediatric cancer groups in %EM or TCR diversities (FigSB-D), the latter suggesting there was no detectable clonally expanded T-cell population in these cancer groups at baseline. Our findings collectively reveal co-elevation of immune checkpoint proteins and central memory T cells subsets in lymphomas at baseline while no evidence of clonal proliferation was observed, in alignment with known knowledge of PD1 regulation of memory T-cell formation ^23–25^.

In pre-treatment solid tumors, we found a trend toward higher levels of naïve T cells compared to patients with leukemias (medians, 35.7% and 21.2%, two-sided Kolmogorov–Smirnov test, p = 0.06, Fig1G). This was accompanied by significantly higher levels of CM% compared to leukemia (medians, 8.2% and 2%, two-sided Kolmogorov–Smirnov test, p = 0.002, Fig1F). We found significantly higher TE% in leukemia compared to solid tumors (medians, 72.4 and 34.5, two-sided Kolmogorov–Smirnov test, p = 0.009, Fig1H). We did not observe any significant difference among SCM% and EM%, nor among TCR and cfTCR diversity across pediatric cancers (FigS2A-D). These results reveal less differentiated T-cell populations in solid tumors compared to leukemia suggesting low antigen exposure.

### Over the course of standard of care, peripheral T cells increase expression of immune checkpoint proteins accompanied by elevated levels of central memory T cells

We next asked how peripheral T cells were affected by standard therapy and whether they are associated with relapsed disease. Here, we compared changes in T-cell subsets and TCR repertoires in post-therapy samples compared to the pre-therapy counterparts.

In leukemia, T-cells collected prior to chemotherapy cycle 4 (pre-cycle 4) showed a significant increase in CM% compared to pre-cycle 1 (median, 0.43 scaled change relative to pre-cycle 1, two-sided t-test with Dunnett’s correction, p = 0.05, FigS3A), consistent with previous report showing an increase in central memory T cells in bone marrow post-chemotherapy in AML^26^. Within leukemia samples, ALL cases showed post-therapy changes in naïve T cells within the 10th-90th percentile range (−1.98, 1.6), whereas two cases with HR ALL showed decrease less than 10th percentile of changes (−2.59 and −3.3 at pre-cycle 3) across all leukemia cases, one of which had a relapsed disease (Fig2A). Two cases with ALL and relapsed HR ALL also showed outlier positive change in CM% over 90th percentile (> 1.96) of leukemia cases (3.83 at pre-cycle 4 and 4.12 at pre-cycle 3, Fig2A). PD1% and TIM3% showed an increasing trend toward pre-cycle 4 compared to pre-cycle 1 (medians, 0.59 for PD1% and 0.58 for TIM3%, two-sided t-test with Dunnett’s correction, p = 0.06 and 0.06, FigS3B). While there was no significant change in cfTCR diversity post-therapy, we observed three ALL cases decreased in cfTCR diversity less than 10th percentile of the leukemia group (< −1.05), while one relapsed AML, and one HR ALL showed increase in cfTCR diversity more than 90th percentile (> 2.11) (Fig2B). This analysis suggests increasing levels of immune checkpoint proteins in leukemia as a result of exposure to chemotherapy.

**Figure 2.**
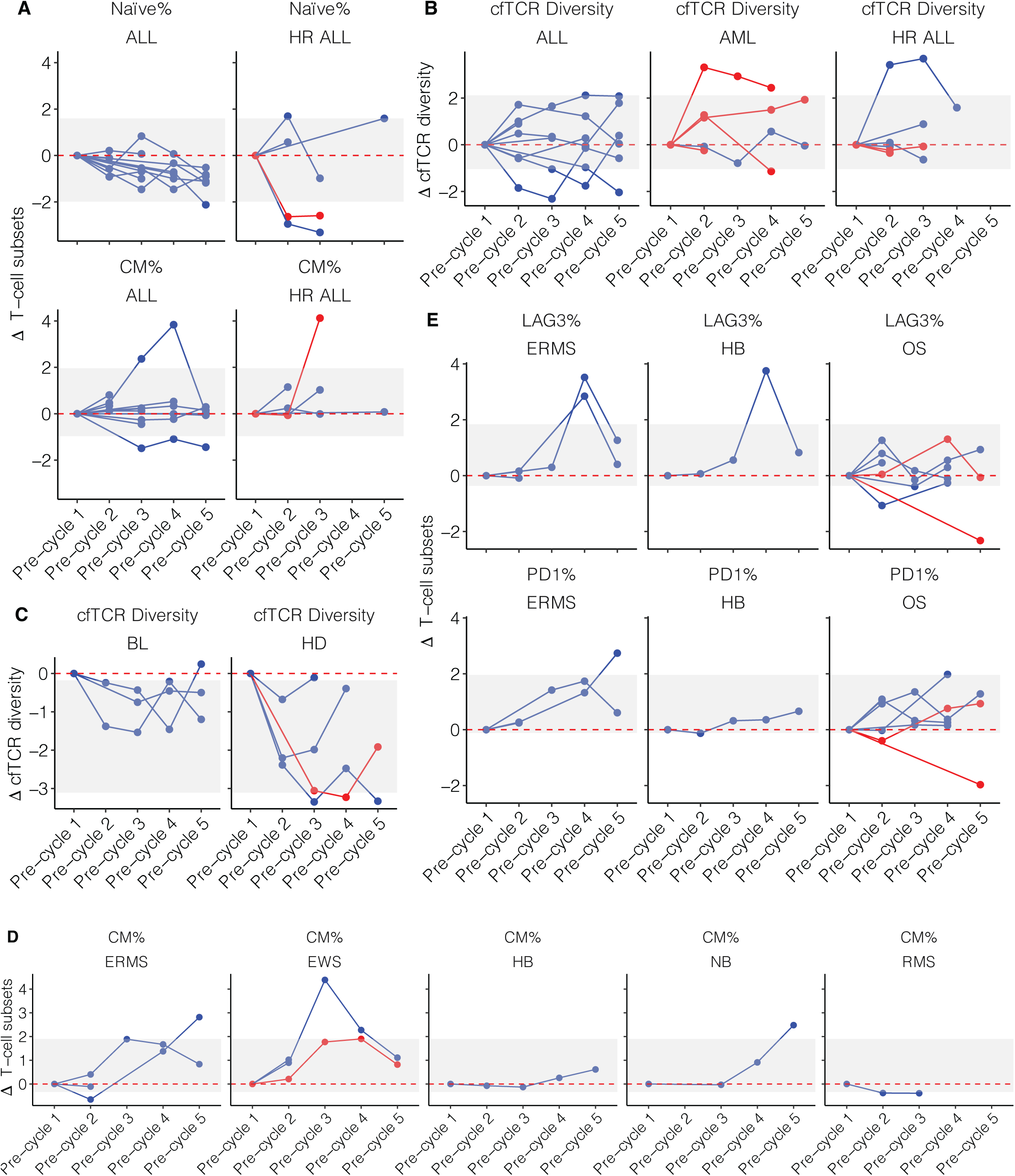
Patient-specific changes in functional and repertoire profiles identifies general and case-specific patterns. **A)** Spider plots depicting patient-specific changes in Naïve% (top panel) and CM% (bottom panel) in ALL (left panel) and HR ALL (right panel) over five cycles of therapy. **B)** Spider plots depicting patient-specific changes in cfTCR diversity in ALL (left panel), AML (middle panel), and HR ALL (right panel) over five cycles of therapy. **C)** Spider plots depicting patient-specific changes in cfTCR diversity in BL (left panel) and HD (right panel) over five cycles of therapy. **D)** Spider plots depicting patient-specific changes in CM% in ERMS, EWS, HB, NB, and RMS (respectively from left to right) over five cycles of therapy. **E)** Spider plots depicting patient-specific changes in LAG3% (top panels) and PD1% (bottom panels) in ERMS (left panels), HB (middle panels), and OS (right panels) over five cycles of therapy. In all plots, each line corresponds to one patient and y-axes show scaled change relative to pre-therapy samples. Grey area denotes 10^th^ – 90^th^ percentile range for each measurement. Red color denotes relapsed disease, while blue color shows patients with complete remission.

In lymphoma, naïve% T cells decreased significantly in pre-cycle 2 and 3 compared to pre-cycle 1 (medians −1.43 and −1.27 relative scaled change, two-sided t-test with Dunnett’s correction, p = 0.02 and 0.007, FigS3D). Comparing case-specific patterns, we observed one case with Burkitt’s lymphoma showing decrease in CM% accompanied by decrease in immune checkpoint proteins (FigS3D). While there was no meaningful change in TCR diversity, cfTCR diversity decreased significantly in all post-therapy samples (medians −0.87, −1.1, −1.45, and −1.19, for pre-cycles 2, 3, 4, and 5, respectively, two-sided t-test with Dunnett’s correction, p = 0.03, 0.001, 0.009, and 0.04, respectively, FigS3F). Three cases with Hodgkin lymphoma showed a decrease of less than 10th percentile (< −3.1) in cfTCR diversity, while cases Burkitt’s lymphoma showed the least change relative to their baseline (Fig2C). These results show that in contrast to leukemia, naïve T cells were most affected by chemotherapy in lymphoma.

In solid tumors, the CM% increased significantly in each post-therapy time point for up to 4 cycles compared to pre-treatment samples (medians 0.07, −0.01, and 0.08, for pre-cycle 2, 3, and 4, respectively, two-sided t-test with Dunnett’s correction, p = 0.03, 0.02, and 0.001 respectively, FigS3G). Three cases with ERMS, EWS, and NB had increase more than 90th percentile (> 1.9) across solid tumors, while HB and RMS remained stable with maximum of 0.6 change relative to pre-therapy (Fig2D). PD1% increased significantly in samples collected pre-cycle 3, 4, and 5 compared to pre-cycle 1 (medians, 1.35, 1.24, and 0.77, respectively, two-sided t-test with Dunnett’s correction, p = 0.008, 0.003, and 0.02, FigS3H). Similarly, TIM3% increased significantly post-therapy (medians, 1.33, 1.28, and 0.76, respectively, two-sided t-test with Dunnett’s correction, p = 0.008, 0.004, and 0.02, FigS3H). LAG3% increased in pre-cycle 4 (median, 0.97, two-sided t-test with Dunnett’s correction, p = 6.6 x e-4, FigS3H). We noted that in an infantile hepatoblastoma case, levels of PD1, and TIM3 were stable post-therapy, while LAG3 increased dramatically in pre-cycle 4 (Fig2E). Similarly, two cases with ERMS showed an increase in LAG3 in pre-cycle 4 (Fig2E). Across six OS cases, two cases with relapsed disease showed a decrease of less than 10th percentile (< −0.11) of solid tumors in levels of PD1% at pre-cycle 5 and 2 (Fig2E). While associations with relapse were limited to small sample size, our results reveal that compared to leukemia and lymphoma, expression of immune checkpoint proteins is elevated post-chemotherapy in solid tumors.

In aggregate, while our study is limited by sample size across cancer entities, our findings reveal an overall pattern of increasing immune checkpoint proteins LAG3, TIM3, and PD1 as a result of chemotherapy, most prominently observed in solid tumors followed by leukemia. Peripheral T-cell composition was affected in a cancer-specific and case-specific manner, with decreasing levels of naïve T cells in lymphoma and increasing levels of central memory cells in solid tumors and leukemia.

### Longitudinal immunogenomic profiling suggests associations with relapsed disease

Having observed notable shifts in peripheral T-cell subsets, immune checkpoint proteins, as well as TCR repertoire in relapsed cases, we next sought to systematically investigate the associations of immunogenomic properties with clinical outcome. Baseline measurements across 11 features (Absolute T-cell Count (ATC), Naïve%, SCM%, CM%, EM%, TE%, PD1%, TIM3%, LAG3%, TCR diversity, and cfTCR diversity) showed no significant associations with relapse (False Discovery Rate (FDR) > 0.2, FigS4A, TableS2). We then calculated patient-specific changes over time for each of these biomarkers using quantile linear regression for each feature in more than two post-therapy samples. We observed no significant associations between longitudinal immune features and incidence of relapse (FDR > 0.2, FigS4B, TableS3).

Having ruled out any single biomarker as a predictor of relapse, we next sought to incorporate functional and repertoire features to quantify overall changes in T-cell population as a result of standard therapy. Principal Component Analysis (PCA) incorporating 11 immunogenomic features across all timepoints demonstrated that the first principal component accounted for 31.3% of the total variance, while the second principal component explained an additional 19.8% (Fig3A). The first principal component (PC) was associated with levels of LAG3%, TIM3%, PD1%, and CM%, so we subsequently termed this the Inverted Immune Checkpoint axis. The second PC had the greatest weights ascribed to Naïve%, TCR and cfTCR diversity, and SCM% (Fig3A), so we termed this the T-Cell Diversity axis. Notably, the terminally differentiated T-cell subset (TE%) demonstrated an inverse association with other variables on both axes, aligning with its biological characteristics of a low proliferation rate and high expression of immune checkpoint proteins. To adjust for potential confounders, we applied a linear mixed-effect model adjusting for individual variations, age, and cancer groups. However, this did not uncover any significant association between changes in PCs and treatment cycles. Patient-specific dynamics of both PCs during standard therapy identified two outlier cases: both with relapsed AML (CHP_413 and CHP_353). While 13 patients displayed changes throughout their treatment ranging from −2.9 to 2.9, these two relapsed cases showed positive change in the Diversity axis (Fig3B). These results revealed two major functional and repertoire immune axes in pediatric cancer patients, while suggesting they may be associated with relapsed disease.

**Figure 3.**
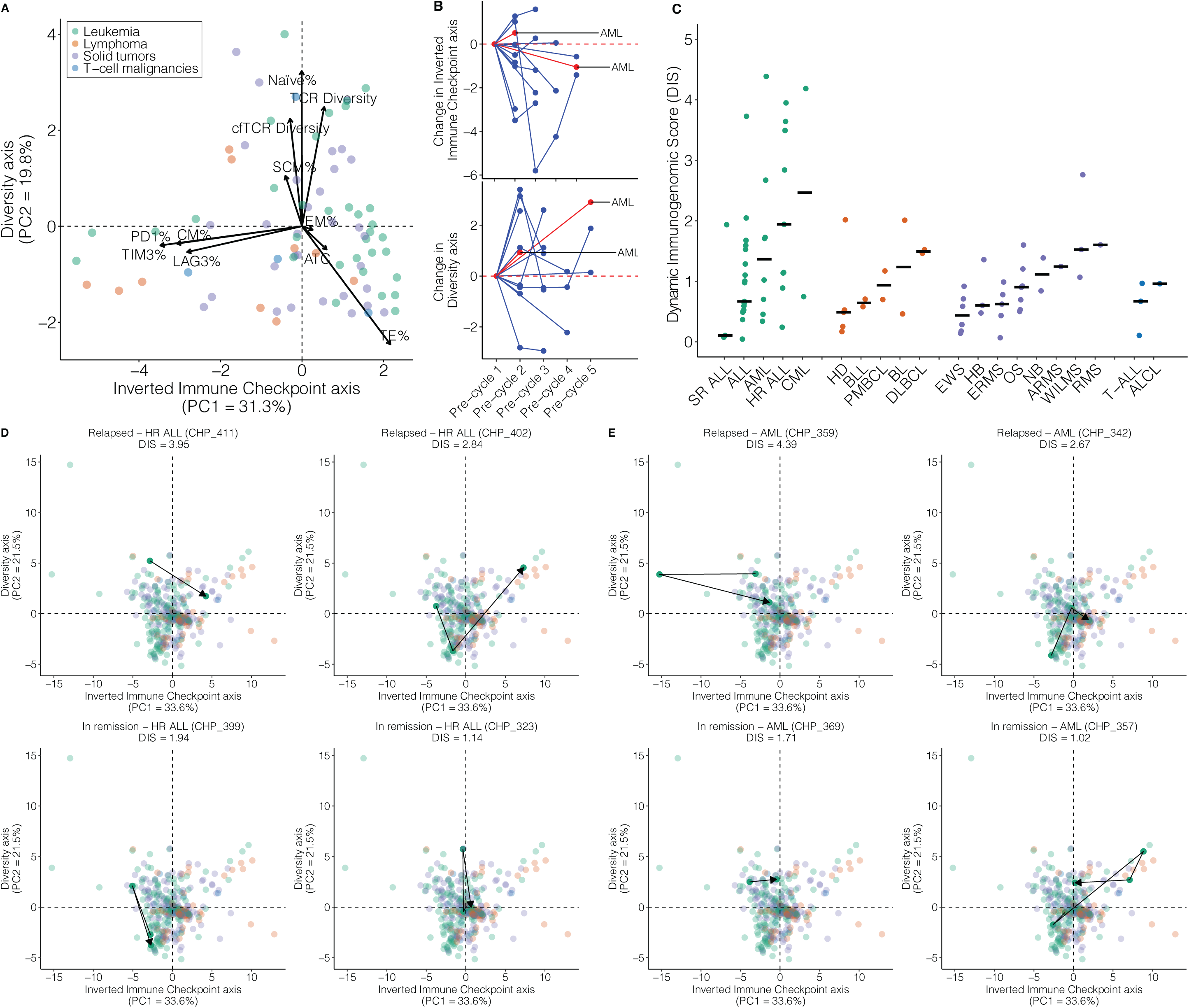
Dynamic Immunogenomic Score (DIS) is associated with high risk cancers and relapsed disease. **A)** PCA of all samples across all functional and repertoire features, colored by cancer group. Arrows show contribution of annotated feature to the PC1 and PC2. **B)** Spider plots depicting patient-specific changes in the Inverted Immune Checkpoint axis (PC1, top panel) and Diversity axis (PC2, bottom panel) over time relative to pre-treatment samples over five cycles of therapy. Each line corresponds to one patient and y-axes show scaled change relative to pre-therapy samples. Red color denotes relapsed disease, while blue color shows patients with complete remission. Cancer types for relapsed cases are annotated on the right. **C)** Breakdown of DIS (y-axis) by cancer type (x-axis) based on imputed PCA. Cancer types are ordered by their median (black bar) within their cancer groups. **D-E)** Immunogenomic trajectories during standard therapy for HR ALL (D) and AML (E) exemplar cases within the imputed PCA space. Arrows connect sequential samples for each case. The DIS on top of each panel is calculated as the median of Euclidean distances between each sequential sample normalized by the number of samples.

To quantify dynamic changes across both axes, we calculated distance metrics for each pair of sequential samples within the two-dimensional PCA space. We considered the median of differences between each timepoint divided by the number of samples as the patient’s “Dynamic Immunogenomic Score (DIS)”. High DIS indicates high variability in the peripheral T cells as patients go through their standard therapy (range: 0.51-1.96, median 0.85). Only 13 patients in our dataset had complete immune profiling with at least one pair of sequential samples, limiting our ability to analyze associations with relapsed disease. To address this limitation, we used the missMDA method ^27^, which imputes missing data points by leveraging the original PCA while accounting for relationships among individuals and variables. Within this imputed dataset, cases with standard risk ALL showed low median DIS compared to high-risk ALL (medians, 0.1 and 1.9, Fig3C). In lymphomas, HD had the lowest median and the more aggressive Burkitts and DLBCLs had the greatest DIS across lymphoma cases (medians, 0.49, 1.23, and 1.49, respectively, Fig3C). In solid tumors, one of the most benign tumors, hepatoblastoma (HB), showed low median DIS compared to WT and RMS (medians, 0.6, 1.52, and 1.6, respectively, Fig3C). We found a significant association with the relapsed disease as univariate (OR: 1.8, CI: 1.1-3.1, Wald test, p = 0.02, TableS4), and particularly in the leukemia patients (including interaction term with cancer groups, OR: 2.4, CI: 1.24-4.65, Wald test, p = 0.009, TableS4). Of note, two cases with relapsed HR ALL that remained in remission upon therapy with Kymriah harbored high DIS (2.84 and 3.9 for patients CHP_411 and CHP_402, Fig3D). Conversely, two HR ALL cases with no incidence of relapsed showed low DIS (1.94 and 1.14 for CHP_399 and CHP_323, Fig3D). Similarly, two relapsed AML cases (CHP_359 and CHP_342) showed higher DIS compared to no relapsed counterparts (CHP_369 and CHP_357) (4.39 and 2.67, 1.71 and 1.02, Fig3E). While further validation is warranted to confirm its predictive value, our results demonstrate that dynamics of immunogenomic states during first-line therapy may be an early indicator of higher risk of relapse.

### TCR repertoire analysis reveals post-therapy increase in TCR specificity groups associated with human and viral antigens in leukemia

Building on the DIS that indicates the treatment’s impact on the overall immune system, we next sought to determine the treatment’s effects on the T-cell repertoire. Using CapTCR-seq applied to 197 PBMC and 266 cfDNA samples from 99 patients, we identified 53,686 TCR sequences. To identify similarities among T-cell repertoires, we calculated pairwise percentage overlap between each two pairs of samples within our cohort. There was no detectable overlap among patients pre- and post-chemotherapy samples indicating the lymphodepleting effects of chemotherapy. Hierarchical clustering of cfDNA samples based on their repertoire overlap identified 18 samples spanning across therapy cycles and cancer types with percentage overlap ranging from 3% to 88.7% (median, 22.2%, Fig4A-B). We further investigated recurring clonotypes within our dataset in further detail. Out of 50,514 unique sequences, 1,482 (2.9%) unique sequences were public, i.e. occurring in at least one sample in more than one patient (range: 2-27 patients) and 1690 were occurring in more than one sample within patients. We used ERGO-II (pEptide tcR matchinG predictiOn) ^28^ to predict peptide-TCR binding of the public TCR sequences. This analysis revealed 17 TCR sequences with predicted binding to peptides originating from human (INS, MLANA), CMV (IE1, pp65), and EBV (BMLF1) (TableS5) suggesting these sequences may recognize common pathogens and self-antigens.

**Figure 4.**
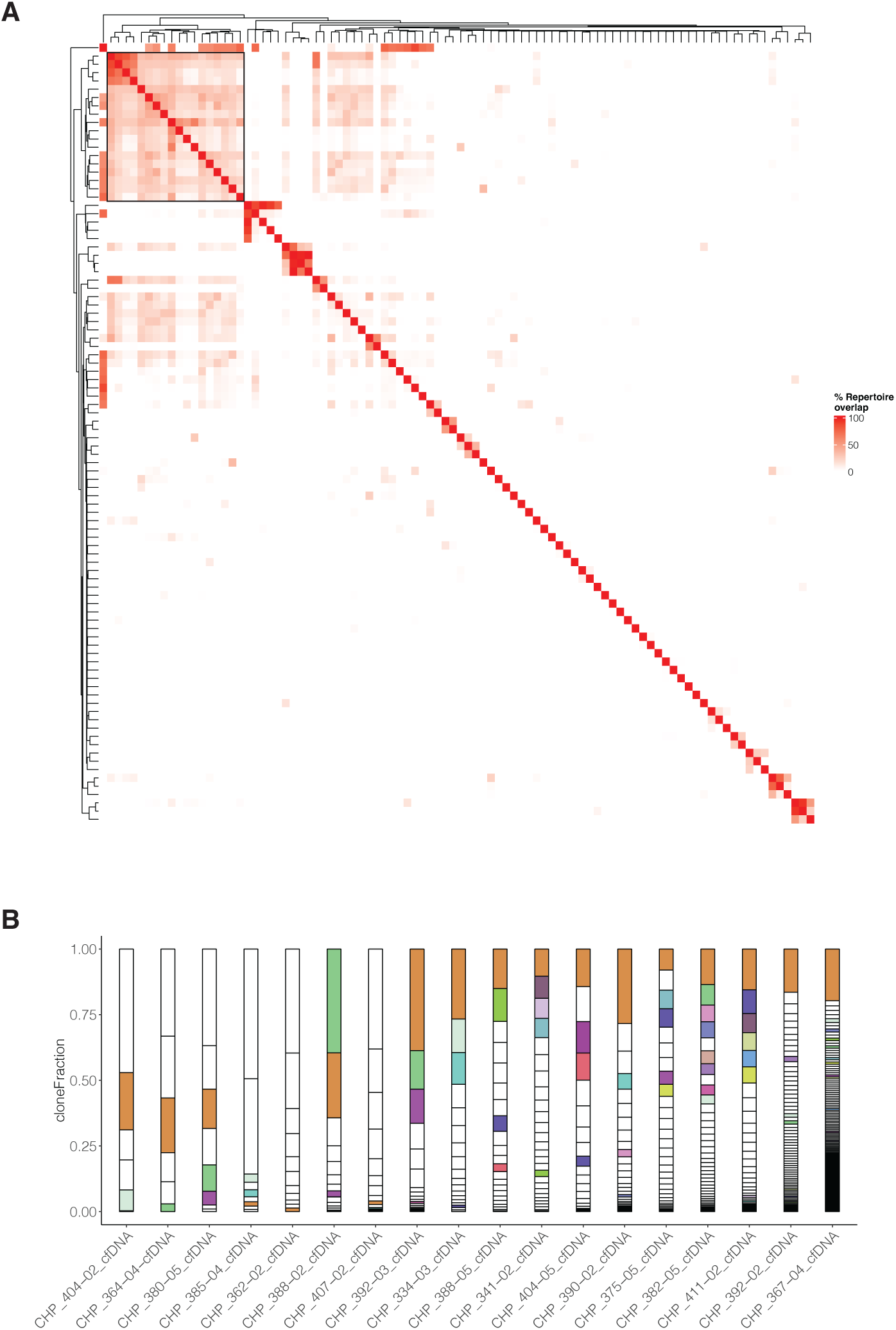
cfTCR analysis identifies overlap in TCR profiles with predicted reactivity to viral and human antigens. **A)** Heatmap depicting hierarchical clustering of repertoire overlap across all cfDNA samples. **B)** Stacked barplot showing clonal fraction (y-axis) of 18 highly overlapped repertoires (x-axis) from A (black box). Colors indicate shared clonotypes across at least two samples.

We next sought to infer the antigen specificities for TCR sequences identified in pediatric cancers. We applied the GLIPHII (Grouping of lymphocyte interactions by paratope hotspots) algorithm^29^ that clusters TCR sequences based on their CDR3β similarities and compares them to a naïve T-cell reference set. We first asked whether there was a systematic difference among adult and pediatric cancers. To address this, we performed GLIPHII analysis using a publicly available set of 25,566 CDR3β sequences collected from adult and pediatric AML patients (NCI TARGET) ^30^. Forty three percent (1,107/2,535) of GLIPHII specificity clusters within pediatric AML samples overlapped with 27% (1,107/4,148) of specificity clusters within adult AML samples (FigS5A). Pediatric AML patients harbored a significantly lower fraction of unique CDR3βs per sample forming a specificity group compared to their adult counterparts (medians 0.41 and 0.5 per patient in pediatric and adult AML, two-sided Student’s t-test, p = 1.6e-5, FigS5B). Across all patients, there was a weak but significant correlation between the fraction of unique CDR3βs in specificity groups and patient age at diagnosis (Pearson’s correlation coefficient = 0.24, p = 0.001, FigS5C). These findings suggest reduced TCR reactivity in pediatric AML patients compared to adults, potentially due to limited antigen exposure.

To assess whether our findings were influenced by the GLIPHII reference TCR dataset, which was derived from adult healthy donors^29^, we repeated our analysis using a custom pediatric-specific TCR reference set. This dataset comprised 9,070,219 CDR3β sequences obtained from 94 samples collected from 25 healthy children (age 1-19 years)^31^. This analysis recapitulates our findings with the default GLIPHII reference set (FigS5A-B). As there were TCR specificities derived from default and pediatric reference sets varied based on distributions of their Fisher scores (median Fisher score, 0.007 and 0.001, FigS5D), we compared all our subsequent analyses with the TCR reference set of pediatric healthy subjects.

We sought to determine the effects of therapy on frequencies of TCR specificity groups in our primary cohort. We chose Simpson diversity of TCR specificity groups to incorporate clone counts and determine changes in abundant groups. In four lymphoma patients, Simpson diversity of TCR specificity groups decreased in post-therapy samples compared to baseline (FigS6A). Among 15 leukemia patients analyzed, 3 of 4 AML cases and 2 of 6 ALL showed similar decreasing trends, whereas both CML cases and three HR ALL cases exhibited increase in Simpson diversity post-therapy compared to baseline levels. In the 12 solid tumor cases, we observed decrease in Simpson diversity in 2 OS, 2 EWS, 1 HB, and 1 NB case. Although these trends did not achieve statistical significance, these trends suggest that decreased diversity may reflect chemotherapy-induced immunogenic cell death, while increased diversity could indicate lymphodepleting effects of therapy.

To annotate TCR specificity groups with potential cognate antigens, we incorporated sequences of known epitope specificities from publicly available databases and published studies^32^ into our GLIPHII analysis^33^. Among the total 21,321 TCR specificity groups identified in 98 patients, 636, 83, and 434 groups were associated with viral (CMV, EBV, HPV, HCV, MCPyV, and Influenza), bacterial (M.tuberculosis and S-pneumoniae), and human epitopes, respectively (TableS6). To investigate differences in the clonal fraction of TCR sequences associated with specific antigens between pre-therapy samples and healthy individuals, we conducted GLIPHII analysis using TCR sequences from 253 healthy adults (age >19 years) and 21 children (age ≤19 years)^29^. To ensure comparable sequencing depth across datasets, we randomly sampled clones based on distribution of total TCR sequences observed in our dataset (Methods, FigS7) and performed analysis of covariance (ANCOVA) to adjust for age. Pre-therapy leukemia samples showed a trend toward lower clonal fraction of TCRs associated with human antigens compared to healthy subjects (medians, 0.01 and 0.02, two-sided Wald test, p = 0.07, Fig5A). We observed significantly lower HPV-associated clonal fraction in pre-therapy samples collected from solid tumor patients compared to adult healthy controls (two-sided Wald test, p = 0.001, Fig5A). These associations suggest low TCR reactivity to common antigens in pre-therapy cancer samples compared to healthy controls.

**Figure 5.**
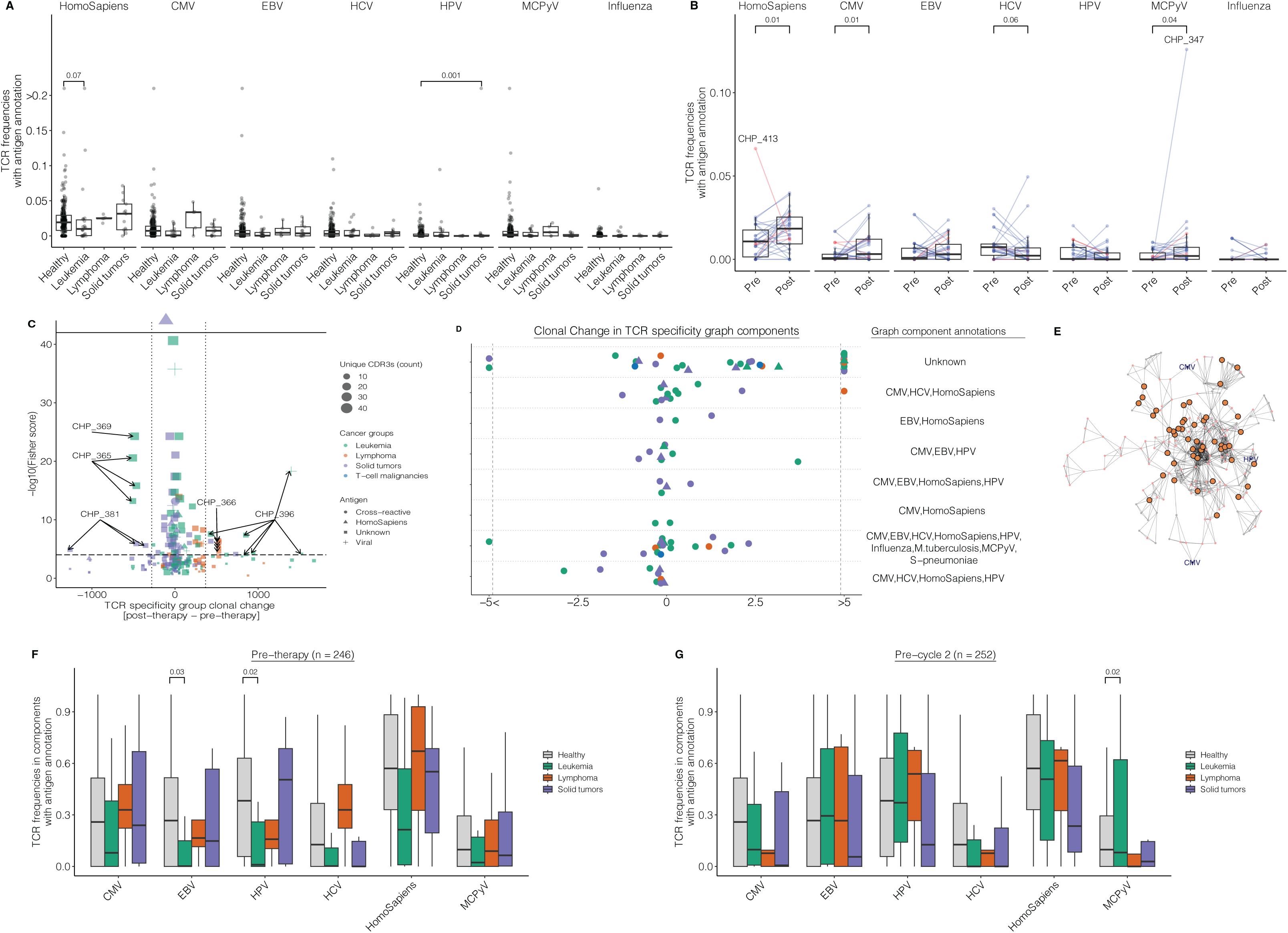
Antigen specificity inference reveals low reactivity in pre-therapy that increases post-therapy in leukemia. **A)** Boxplot showing frequency of TCRs within specificity groups that were annotated with human or viral antigens across pre-therapy pediatric cancer groups and healthy subjects (ANCOVA adjusting for age, two-sided Wald test). **B)** Boxplot showing frequency of TCRs within specificity groups that were annotated with human or viral antigens across paired pre- and post-therapy leukemia samples (Wilcoxon signed-rank test). **C)** Scatterplot illustrating clonal change (x-axis) of significant TCR specificity groups, as determined by Fisher’s exact test (y-axis, −log scale), shared among post- vs pre- therapy samples. Shapes denote antigens and colors show cancer groups. Horizontal dashed line corresponds to Fisher score of 0.0001. Vertical dotted lines correspond to 10^th^ and 90^th^ percentile of clonal change (−280.8 and 367.6). Outlier cases are annotated. **D)** Dot plot showing scaled clonal change (x-axis) of network components (y-axis) shared between pre-therapy and at least one post-therapy sample and occurring in more than one patient. Colors indicate cancer groups. Shapes denote disease status with triangles indicating relapsed disease. Table on the right shows antigen annotations. Scaled clonal changes more than absolute value of 5, were capped (vertical dashed lines) for visualization. **E)** Exemplar network graph showing component 5 consisting of TCRs from pediatric cancer samples (large dots) and healthy subjects (small dots). Antigen annotations are labeled. **F-G)** Grouped boxplots showing TCR frequencies (y-axis) associated with antigens (x-axis) in healthy and pre-therapy (F) or post-therapy (G) samples (ANCOVA adjusting for age, two-sided Wald test). In all boxplots, boxes show median and interquartile range (IQR) and whiskers represent 1.5 × IQR.

We next compared clonal changes of TCR sequences annotated with known antigens among pre- and all post-therapy samples in a pair-wise manner. In leukemia, we found significant increase in clonal fraction of TCR clones associated with human, CMV, and MCPyV epitopes in post-therapy samples compared to pre-therapy counterparts (medians, 0.018 & 0.011, 0.003 & 0.0007, 0.002 & 0, for TCR clones associated with human, CMV, and MCPyV, respectively, Wilcoxon signed-rank test, unadjusted p = 0.01, 0.01 & 0.04, FDR < 10%, Fig5B). Among the three relapsed cases, one AML case (CHP_413) exhibited an opposite pattern, with T-cell clones associated with melanoma-associated antigen (MAA)^35^ decreasing post-therapy (clonal fraction 0.06 and 0.01 for pre- and post-therapy, Fig5B). Conversely, HCV associated clones decreased in post-therapy samples (medians 0.002 & 0.007, Wilcoxon signed-rank test, unadjusted p = 0.06, FDR < 10%, Fig5B).

In lymphoma, we did not detect any significant changes between pre- and post-therapy sample pairs. However, with the exception of an HD case (CHP_415), TCR sequences associated with human epitopes showed a decreasing trend post-therapy compared to pre-therapy (median clonal fraction 0.02 and 0.005 for pre- and post-therapy, FigS6B). Similarly, T-cell clones associated with CMV epitopes, with the exception of another HD case (CHP_366), exhibited higher clonal fractions in pre-therapy samples (medians clonal fraction 0.03 & 0 for pre- and post-therapy, FigS6B).

In solid tumors, while there were no significant changes in clonal fraction between pre- and post-therapy, TCR sequences associated with human and CMV epitopes exhibited a similar decreasing trend as lymphoma post-therapy compared to pre-therapy (median clonal fraction 0.03 and 0.02 for human epitopes, 0.01 and 0.005 for CMV epitopes, pre- and post-therapy, FigS6C). Among the relapsed cases, two relapsed cases (CHP_418 with NB, CHP_364 with OS) showed high fraction of clones associated with EBV (clonal fractions, 0.01 and 0.03) that were further increased in post-therapy samples (clonal fraction, 0.05 and 0.04, FigS6C). Together, our findings highlight an increase in T-cell reactivity to human and CMV epitopes following chemotherapy in leukemia, contrasted by opposing trends observed in lymphoma and solid tumors.

We further explored patient-specific clonal dynamics of TCR specificity groups detectable in pre- and post-therapy samples. In 101 paired pre- and post-therapy PBMC samples from 33 patients, 245 TCR specificity groups were present in at least one paired sample set from a patient. Seven groups showed reduction less than 10th percentile (< −280.8) in post-therapy samples relative to pre-therapy. Fourteen groups showed expansion more than 90th percentile (> 367.6) in post-therapy samples relative to pre-therapy (Fig5C). A one-year old HR ALL patient (CHP_396) exhibited dramatic clonal expansion of a TCR specificity group “SP%RNTE’’ that was associated with a CMV pp65 epitope. The same patient showed post-therapy clonal expansion of one TCR specificity group (> 1500, SPPTGDELL%KNI) and four other groups with unknown specificities, with clonal expansions ranging from 431 to 941 clones. We speculate that chemotherapy may have triggered CMV reactivation in this patient, contributing to the observed clonal expansion. Conversely, an ALL patient (CHP_365) and an EWS case (CHP_381) exhibited three TCR specificity groups that underwent significant clonal shrinkage post-therapy, falling below the 10th percentile for clonal change (Fig5C). While experimental validation is warranted, our findings reveal patient-specific patterns of T-cell reactivity in pediatric cancers in response to standard therapy.

We noted that TCR sequences may form multiple specificity groups. We therefore performed network analysis of TCR sequences that share TCR specificity groups^33^. We grouped together specificity groups that share identical CDR3β clones into “components”. Among the 33 patients with pre-therapy and at least one post-therapy PBMC sample, we found a total of 22 components, eight of which occurred in more than one patient (Fig5D, TableS7). We observed that 7 of these 8 components were annotated with viral antigens, with an average change of 1,650 clonal fraction (median change 78 relative to pre-therapy) (Fig5D). To determine whether this bias towards viral specificity was specific to patients with pediatric cancers, we performed an additional analysis leveraging publicly available pediatric and adult healthy subjects (Methods). In this analysis, we found 199 components spanning healthy and pre-therapy samples. Pre-therapy leukemia samples harbored significantly lower TCR frequencies in components associated with the EBV and HPV antigens compared to healthy subjects (medians, 0.002 and 0.27 for EBV, 0.009 and 0.38 for HPV, ANCOVA adjusting for age, two-sided Wald test, p = 0.03 and 0.02, Fig5F). Conversely, after one cycle of therapy, we found that TCR clonal fraction within components associated with MCPyV antigens increased after adjusting for age compared to healthy subjects (medians, 0.37 and 0.38, ANCOVA adjusting for age, two-sided Wald test, p = 0.02, Fig5G). However, these associations appeared to be exclusive to leukemia, as we did not find any significant difference among components derived from lymphoma, solid tumors, and healthy cases. While further investigations are warranted to identify disease-specific TCR sequences with their reactive antigens, these associations suggest leukemia patients may be less exposed to common antigens that may in turn contribute to immunosurveillance.

## Discussion

In the current study, we provided multimodal immunogenomic analysis of peripheral T cells in children undergoing standard therapy. We found high levels of immune checkpoint proteins, PD1, LAG3, and TIM3, accompanied by high percentages of central memory T cells in lymphoma patients. Our results were in line with previous reports elucidating that PD1 regulates memory T-cell differentiation ^23,36^. We speculate that targeting the PD1 in lymphoma patients leads to amplification of weak TCR signals and improved anti-tumor immunity. This, coupled with high expression of PD-L1 on lymphoma cells, may be the underlying mechanism of favorable response rate in lymphomas ^13–15^.

Dynamic Immunogenomic Score (DIS), while requiring further validation, provides a multimodal approach to model the impact of chemotherapy on T cells. Although the lymphodepleting effects of chemotherapy are widely known, the accurate quantification and associations with disease outcome remain less characterized. One recent study showed that complete loss of T cells resulted in disease relapse in the B-cell ALL mouse model and that elevated levels of CD4+ and CD8+ T cells post-chemotherapy were associated with improved survival in pediatric ALL patients^37^. Regulatory T cells expand in AML patients after intensive timed chemotherapy, however, the prognostic value was not evaluated ^38^. Another study showed that in AML patients post-chemotherapy gene enrichment scores related to CD8+ naïve and central memory T cells were enriched in responders versus non-responders ^39^. Our study extended these studies through measurement of 11 immunogenomic features at baseline before treatment and at multiple timepoints throughout standard of care. This study highlights the potential of multimodal immune analysis for identifying high-risk populations during the early stages of disease. We speculate that the performance and generalizability of the DIS concept may improve with the collection of high-dimensional data, for instance, single-cell RNA-seq and TCR/BCR-seq performed on FACS-sorted T cells. This approach may provide particular utility in cellular immunotherapy, including CAR T-cell therapy as the properties of autologous T cells are critical in manufacturing a successful CAR product, monitoring T-cell persistence, and predicting treatment outcome.

We explored dynamics of specific T-cell clones over time, and mapping antigen specificities using GLIPHII and network analysis. In a previous study comparing lung cancer and adjacent healthy tissue in adults^40^, this approach revealed T-cell cross-reactivity between human tumor-associated antigens and common viral and bacterial antigens. In our study of pediatric cancer patients, we encountered a similar phenomenon whereby high abundance of virally-associated TCRs in healthy subjects compared to leukemia patients may suggest better immune surveillance. Notably, an increase in the clonal fraction of TCRs related to several antigens in post-therapy samples points to the immunogenic effects of chemotherapy, consistent with reports of immunogenic cell death induced by chemotherapy that canalter T-cell reactivity ^41,42^.

Establishing a reference dataset of reactive TCRs and their cognate antigens specifically for pediatric cancers, could accelerate experimental efforts toward novel immunotherapeutic strategies by identifying tumor-reactive TCRs. To support these efforts, we developed T-cell and ImmunoGlobulin Epitope Receptor DataBase (TIGERdb), which serves as a repository to store and reanalyze T-cell repertoires in future studies as more target antigens for TCRs become available and validated experimentally (https://tigerdb.ca).

Although our study reveals the prognostic value of multimodal immune profiling during standard of care, it is subject to several limitations. Firstly, the diverse nature of our cohort resulted in small sample size for individual cancer types–each likely harboring distinct levels of immunogenicity. This hindered our ability to establish disease-specific patterns and identify cancer-specific T-cell clones. Secondly, the relatively small number of relapsed cases within our dataset limited our ability to validate our statistical analysis. Given the potential of liquid biopsies for early detection and therapy response, as highlighted in recent studies ^43,44^, validating our findings in prospective studies could enable early risk stratification in children through the minimally invasive blood draws. Thirdly, although the DIS offers a framework for assessing the overall effect of standard therapy on peripheral T cells, our analysis could not establish direct correlations between immune checkpoint protein expression, TCR diversity, and functional T-cell memory subsets. This is likely due to the heterogeneous nature of immune responses to pediatric cancer and highlights a need for expanded immunogenomic profiling in childhood cancers. Such correlations could potentially yield a more accurate and predictive model, as well as inform therapeutic and molecular monitoring strategies.

In summary, our findings underscore the value of multimodal immunoprofiling during standard of care to identify patients at higher risk of relapse, enabling personalized clinical management of pediatric cancers.

## Methods

### Participants and sample collection

This study was part of an observational longitudinal study of pediatric patients receiving chemotherapy and was approved by Children’s Hospital of Philadelphia Institutional Review Board (IRB #12-009915) and University Health Network Research Ethics Committee (CAPCR #18-5420). The cohort comprised 119 patients, with ages ranging from 1 month to 21 years, all newly diagnosed with leukemia, lymphoma, or solid tumors, as outlined in Table 1. As of March 2023, 12 patients had died from their disease, and 10 were lost to follow-up. Of the cohort, 23 patients experienced disease relapse, while 96 achieved complete remission. Ten patients (7 AML, 1 CML, 1 HR ALL, 1 HD) received bone marrow transplantation. Peripheral blood samples were collected in Ethylenediaminetetraacetic Acid (EDTA) tubes at diagnosis and subsequently prior to the initiation of each cycle of both induction and maintenance therapy. Plasma was isolated and stored at −80°C. PBMC was isolated using Ficoll-Paque (GE Life Sciences) density centrifugation medium as per standard protocol. The isolated cells were washed in phosphate-buffered saline (PBS) with 2% bovine serum albumin (BSA), counted, and cryopreserved in 90% Fetal Bovine Serum (FBS) and 10% Dimethyl Sulfoxide (DMSO). An aliquot was stained for CD3, CD4, and CD8 expression to obtain ATC.

### Flow cytometry

We performed T-cell phenotyping based on surface expression proteins, as previously described^17,18^. We determined T-cell subsets as follows: Naïve (CCR7^+^, CD62L^+^, CD45RO^−^, and CD95^−^), Stem-like Central Memory (SCM, CCR7^+^, CD62L^+^, CD45RO^−^, and CD95^+^), Central Memory (CM, CCR7^+^, CD62L^+^, CD45RO^+^, and CD95^+^), Effector Memory (EM, CCR7^−^, CD62L^−^, CD45RO^+^, and CD95^+^), and Terminal Effector (TE, CCR7^−^, CD62L^−^, CD45RO^−^, and CD95^+^). The following antibodies were used to characterize and sort T-cell subsets on a BD FACSVerse flow cytometer (BD Biosciences, #653118): CD8–FITC (fluorescein isothiocyanate) (BD Biosciences, #347313), CD3–PE (phycoerythrin) (BD Biosciences, #555340), CD4–APC (allophycocyanin) (BD Biosciences, #555349), CCR7-FITC (BD Biosciences, #561271), CD95-PE (BD Biosciences, #556641), CD45RO (BD Biosciences, #559865), and CD62L-PE/Cy7 (cyanine 7) (BioLegend; 304822). For immune checkpoint proteins, we used the following antibodies: LAG-3 (Invitrogen 46-2239-42), TIM-3 (BD Biosciences, 565562), and PD-1 (BD Biosciences, 563076). Analysis was performed using FlowJo software (Treestar Inc.).

### TCR sequencing and quality control

To study T-cell repertoire in tumor samples, we used hybrid capture TCR sequencing (CapTCR-seq^21^). We extracted a median of 4.3 ug DNA from frozen purified PBMCs and a median of 37 ng cell-free DNA (cfDNA) from plasma using QIAGEN Allprep kit. Genomic DNA was sheared to median 250-bp insert size. Illumina DNA libraries were prepared from a median of 500 ng DNA and 20 ng of cfDNA using the KAPA HyperPrep library preparation kit with custom adaptors, including molecular identifiers ^45^. We performed hybrid capture using a custom TCR probeset^21^, following the Integrated DNA Technologies hybridization capture protocol using 250 ng of indexed libraries. Captured DNA pools were then deep sequenced on the Illumina NovaSeq platform at the Princess Margaret Genomics Centre. We used the MiXCR v.2.1.12 immune repertoire pipeline^46^ to align raw sequencing reads to IMGT reference (repseqio v.1.5), assemble into clonotypes, and export tab delimited files. Of 321 PBMC samples, we successfully prepared 282 libraries, yielding 278 successful captures, sequenced to the median of 5 million reads. Of 316 plasma samples, 316 successful libraries, 295 successful captures, 279 sufficient sequencing reads.

### Dynamic Immunogenomic Score (DIS)

To measure changes over time, we used quantile linear regression scaled to the number of post-therapy samples for each patient using quantreg R package (v5.97). For baseline associations, we used z-scores calculated over all pre-therapy samples. For DIS, we performed dimension reduction on 11 features using the prcomp function in R. For each patient, we calculated the median of Euclidean distances between each subsequent sample scaled to the number of samples for that patient as DIS. Acknowledging missing values in measurements ranging from 1% to 52% of 393 samples, we imputed the missing values using the MissMDA, iterative PCA algorithm^27^ to enable logistic regression analysis predicting incidence of relapse.

### TCR analysis

The repertoire diversity was calculated using the iNEXT v.2.0.20 R package ^48^. The percentage of overlap in TCR repertoire was calculated as in Sun et al. ^45^, where the number of common TCRs found in a pair of samples is divided by the total TCR count for both samples. We used the ERGO-II for peptide binding prediction of public sequences through the TCRosetta platform^49^. The output for ERGO-II analysis is available as TableS5.

### GLIPHII specificity group analysis

The GLIPHII algorithm was used to cluster the CDR3β sequences into specificity groups. Briefly, a GLIPHII pattern is the specific amino acid pattern, either global or local, that defines a cluster; CDR3βs with the same pattern are inferred to have the same specificity. For the external AML samples, the analysis was first done with the default TCR reference provided by GLIPHII, and repeated using the custom pediatric reference set^31^. Briefly, we downloaded the Mitchell el al^31^ dataset from the ImmuneAccess Adaptive portal. The original dataset included TCR-seq data from 216 samples collected longitudinally from 25 healthy children and 29 children with type I diabetes. We removed 13 samples due to productive rearrangements less than 5th of the dataset and created a pediatric reference dataset for GLIPHII analysis using 94 samples collected from 25 healthy children.

For the main specificity analysis, 54,985 CDR3βs from the pediatric cohort were analyzed using GLIPHII, along with 3,681 CDR3βs with known specificities against several types of antigens like viruses and tumor-associated antigens (TIGERdb) and 19,596 CDR3βs from an adult cancer cohort (MDavis) ^40^. Specifically, TIGERdb is a database containing browsable TCRs from over 100 curated publications, with additional information ranging from specificity, HLA-restriction, V- and J-gene usage, and epitope sequence. Antigen specificity can be inferred if TIGERdb TCRs cluster together with TCRs with unknown specificity. The confidence of the inference improves with 1) global patterns, 2) consistent HLA restriction, and 3) similarity in amino acid properties at non-conserved positions in the pattern. The TCRs from the primary cohort are accessible through TIGERdb (https://tigerdb.ca).

We utilized the primary cohort from Emerson et al. (34) as healthy controls. The original dataset comprised TCR-seq data from 666 individuals. We excluded 12 cases with productive rearrangements below the 5th percentile of the dataset (<47,512 rearrangements). Of the remaining cases, 346 were CMV-negative, including 68 without reported age. We constructed a healthy cohort consisting of 21 children (under 19 years) and 256 adults (19 years and older). GLIPHII analysis was performed separately on the adult and pediatric cohorts, and the results were integrated with our primary dataset for further analysis.

### TCR specificity network generation and analysis

We followed Soleimani et al (2024) method for concatenating and pruning the GLIPHII specificity groups via clique-based pruning to form components, and dividing the components into communities based on modularity. Clique-based pruning was done using the max_cliques function in the igraph (v1.5.1) R package, and community detection was done via Leiden clustering using the cluster_leiden function from the same package. Briefly, Leiden clustering is a fast and high-quality clustering approach that maximizes cluster modularity and minimizes connection between different clusters^50^.

To ensure the robustness of the components, we filtered out graphs with fewer than four CDR3βs then used clique-based pruning to remove CDR3βs with low transitivity, as in Soleimani et al. The resulting, fully-connected graph is defined as a component. Finally, we label each component with its corresponding specificity (viral, bacterial, or human). Communities with one of viral or bacterial specificity and *H. sapiens* specificity are considered cross-reactive.

### Statistical analysis

All analyses were performed and plotted on R v4.0 in Jupyter notebooks. We used the two-sided exact Kolmogorov-Smirnov test for comparing differences among pre-therapy samples. To determine feature associations with therapy cycles, we calculated patient-specific differences of z-scores in post-therapy samples relative to pre-therapy. We fitted a linear mixed effect model including age as a fixed effect and patients as a random effect using the lmer function of lme4 R package (v1.1-35.1). We used lsmeans R package (v2.3) to compute and compare least-square means among post- and pre-therapy samples. We used Wilcoxon signed-rank test to compare clonal changes of TCR specificity groups in matched pre- and post-therapy samples. We used two-sided

Fisher’s exact test to compare clonal differences of annotated TCR specificity groups among pediatric and adult AML. We used a two-sided Student’s t-test to compare TCR clonal fractions among pediatric and adult AML patients. We used Pearson correlation to determine the association between age and clonal fraction. To predict incidence of relapse, we fitted a univariate logistic regression model using glm function in R. In all boxplots, boxes show median and IQR, and whiskers represent 1.5 times IQR. Boxplots are shown for groups with more than three data points. Plot aesthetics were edited using Adobe Illustrator v24.0.1.

## Data and Code availability

Clinical and functional data are available as supplementary information to this paper and at https://github.com/pughlab/ped_CapTCRseq. TCR clones are available to query and search at https://tigerdb.ca. CDR3β sequences from the peripheral blood mononuclear cells (PBMCs) of 22 pediatric and 151 adult AML patients from Zhang et al. (2019)^30^ were used to determine whether there was a systematic difference among adult and pediatric leukemias. The pediatric and adult samples were originally from the Therapeutically Applicable Research to Generate Effective Treatments (TARGET) and The Cancer Genome Atlas (TCGA) programs. CDR3β sequences with known antigen specificities used in our analysis are available at https://tigerdb.ca. We also included adult solid cancer CDR3β sequences from Gee et al.^51^ in our GLIPHII analysis. We downloaded reference healthy adult and pediatric controls from Adaptive technologies portal (http://adaptivebiotech.com/pub/mitchell-2022-jcii and http://adaptivebiotech.com/pub/emerson-2017-natgen). Custom codes to reproduce the analyses and figures are publicly available at https://github.com/pughlab/ped_CapTCRseq.

## Acknowledgements

This work was supported primarily by a StandUp2Cancer Phillip A. Sharp Innovation in Collaboration Award to T.J.P. and D.B. Additional funding support for the project was made possible by the Princess Margaret Cancer Foundation, Ontario Institute for Cancer Research and Terry Fox Research Institute. TIGERdb development was supported by a grant from The Leukemia & Lymphoma Society. T.J.P. holds the Canada Research Chair in Translational Genomics and is supported by a Senior Investigator Award from the Ontario Institute for Cancer Research and the Gattuso-Slaight Personalized Cancer Medicine Fund. A.N. was supported by a Princess Margaret Postdoctoral Fellowship. We gratefully acknowledge the individuals from the Princess Margaret Genomics Centre (www.pmgenomics.ca; T. Ketela and J. Tsao) and Bioinformatics Services (C.Virtanen, Z. Lu, and N. Stickle) for their expertise in generating the TCR-sequencing data used in this study. Additional infrastructure support was received from the Canada Foundation for Innovation, Leaders Opportunity Fund (CFI 32383 and 38401) and the Ontario Ministry of Research and Innovation, Ontario Research Fund Small Infrastructure Program. DB is an employee of Kite Pharma, Inc.

## Supplementary Figures

**Figure S1.**
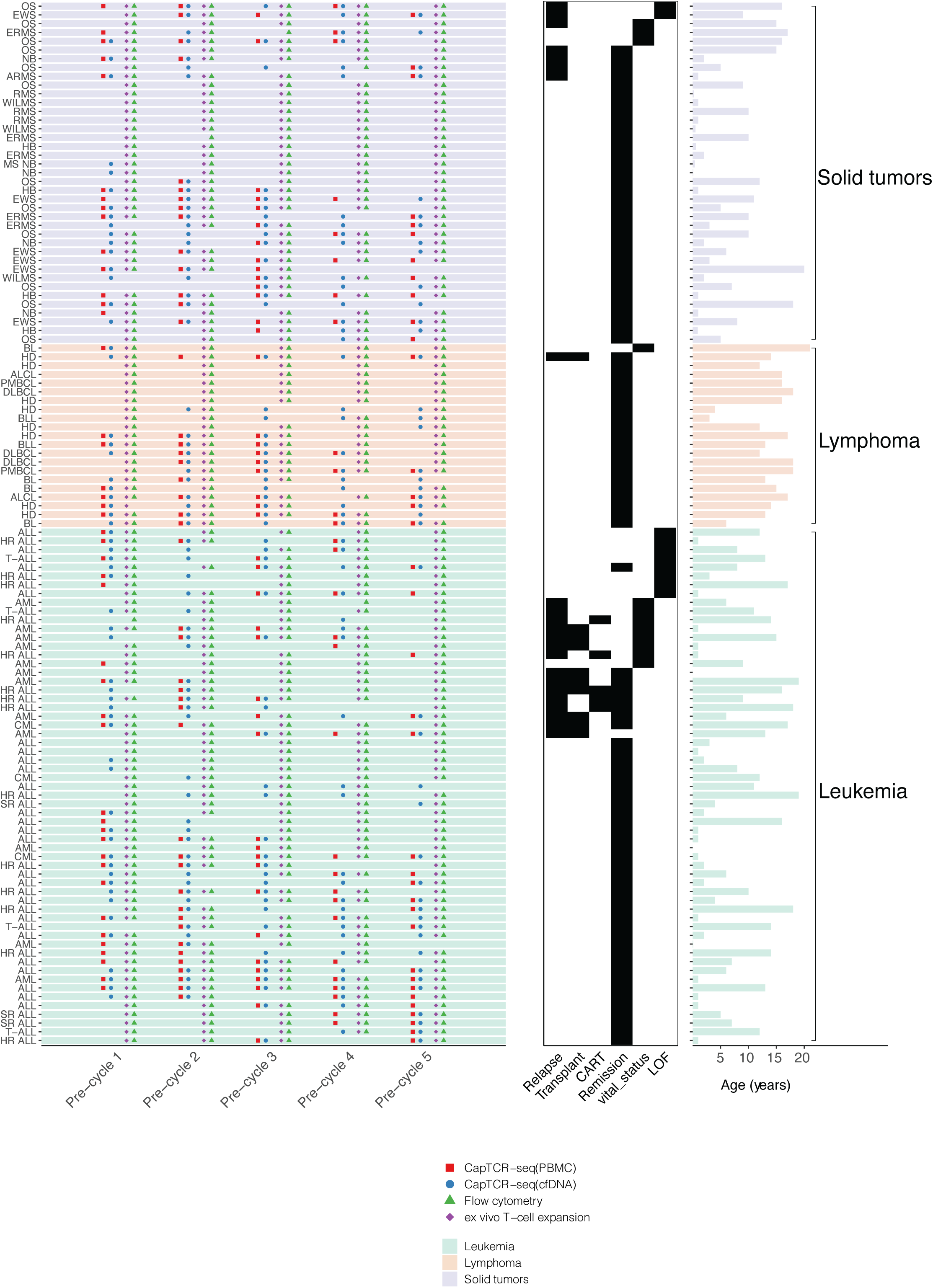
Overview of patient characteristics and assays. The left-hand side plot shows patients (y-axis) and timepoints (x-axis). Each data point indicates the assay performed. Patient’s diagnosis is annotated on the y-axis. Heatmap in the middle shows binary patient outcome (Relapse, Transplant, CAR T-cell therapy, Complete remission, vital status, or loss of follow-up (LOF)). Barplot on the right shows age at diagnosis in years. Plots are ordered by cancer groups (Solid tumors, lymphoma, or leukemia) and patient outcome (heatmap) (N = 119 patients).

**Figure S2.**
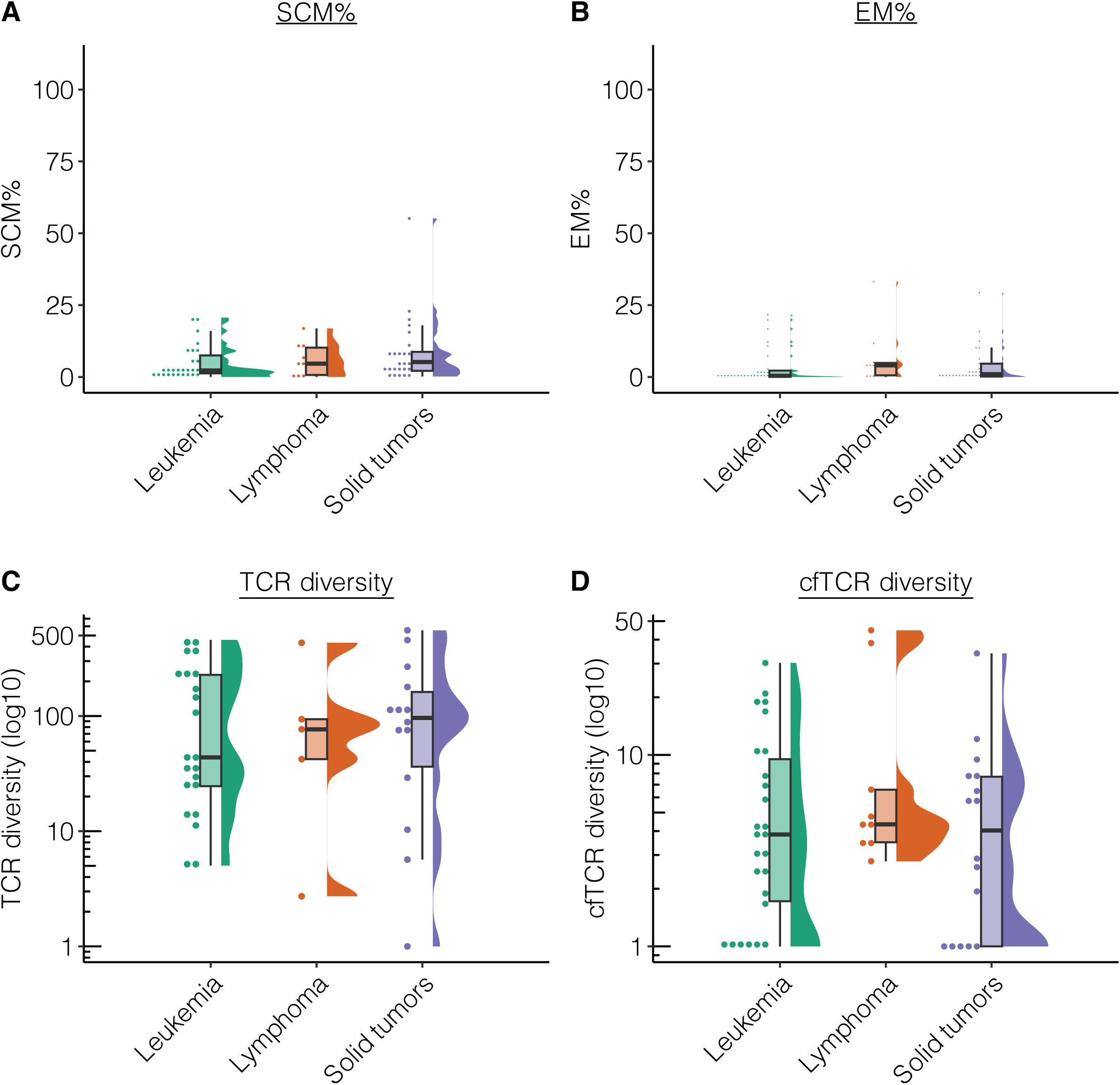
**A-D)** Rain cloud plots showing SCM% (A), EM% (B), T-cell diversity in PBMC (C), and cfDNA (D) pre-therapy across three cancer groups. In boxplots, boxes show median and interquartile range (IQR) and whiskers represent 1.5 × IQR.

**Figure S3.**
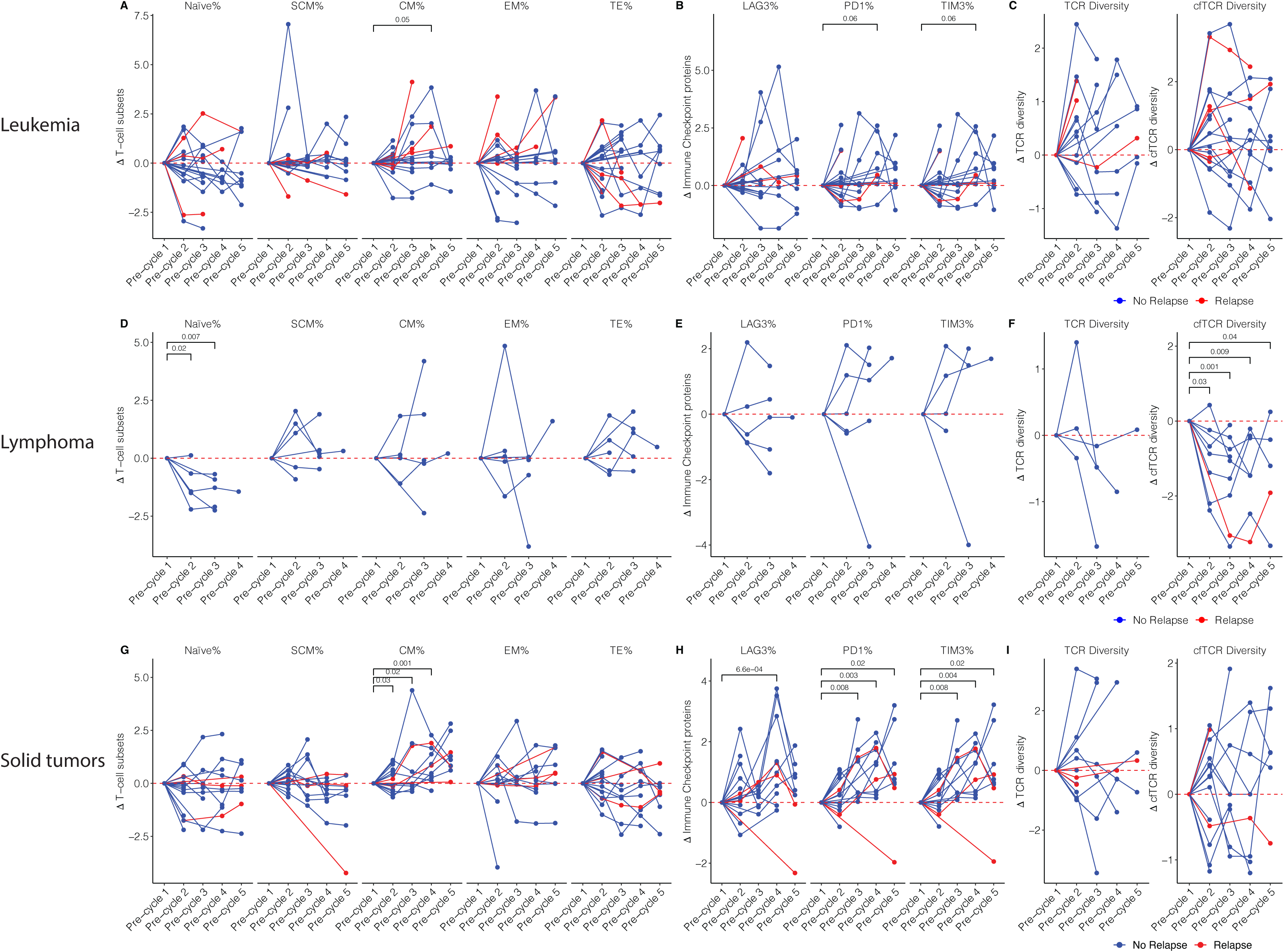
Patient-specific changes in functional and repertoire profiles. **A-I)** Spider plots depicting patient-specific changes in T-cell subsets (A, D, E), Immune checkpoint proteins (B, E, H), and (cf)TCR diversity (C, F, I) over time relative to pre-treatment samples in leukemia, lymphoma, and solid tumors (two-sided t-test with Dunnett’s correction). In all plots, each line corresponds to one patient and y-axes show scaled change relative to pre-therapy samples. Red color denotes relapsed disease, while blue color shows patients with complete remission.

**Figure S4.**
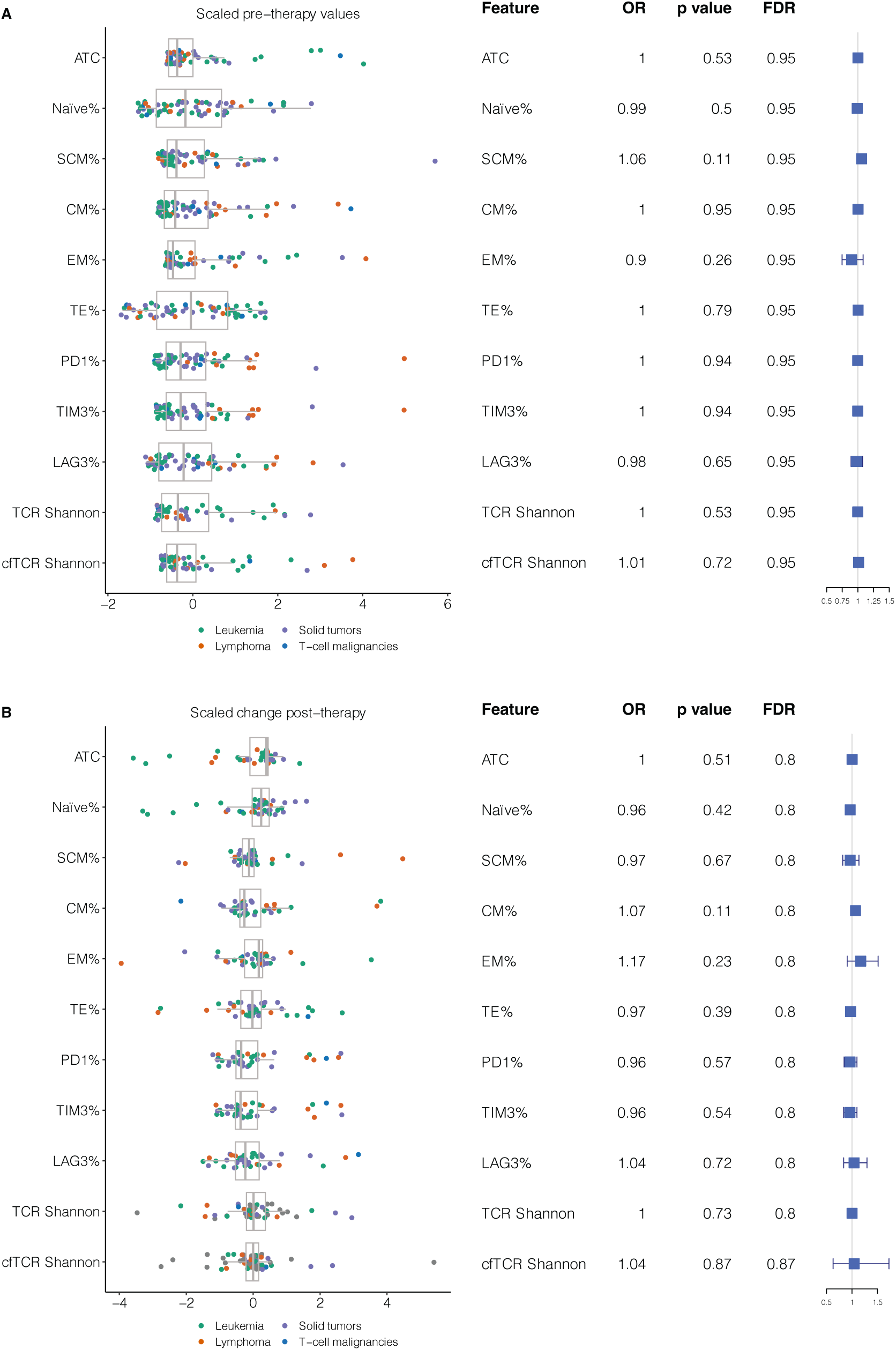
Individual pre- or post-therapy profiling is not associated with incidence of relapse. **A)** Boxplot showing scaled values (x-axis) for each measurement at pre-therapy (y-axis). Each data point is one sample colored by cancer group. Forest plot depicting results of univariate logistic regression analysis for each feature (rows). X-axis shows hazard ratios (boxes) and 95% confidence interval (whiskers) (two-sided Wald test, N = 85 patients). **B)** Boxplot showing scaled change during therapy, calculated using quantile linear regression (x-axis) for each measurement (y-axis). Each data point is one sample colored by cancer group. Forest plot depicting results of univariate logistic regression analysis for each feature (rows). X-axis shows hazard ratios (boxes) and 95% confidence interval (whiskers) (two-sided Wald test, N = 81 patients). In boxplots, boxes show median and interquartile range (IQR) and whiskers represent 1.5 × IQR.

**Figure S5.**
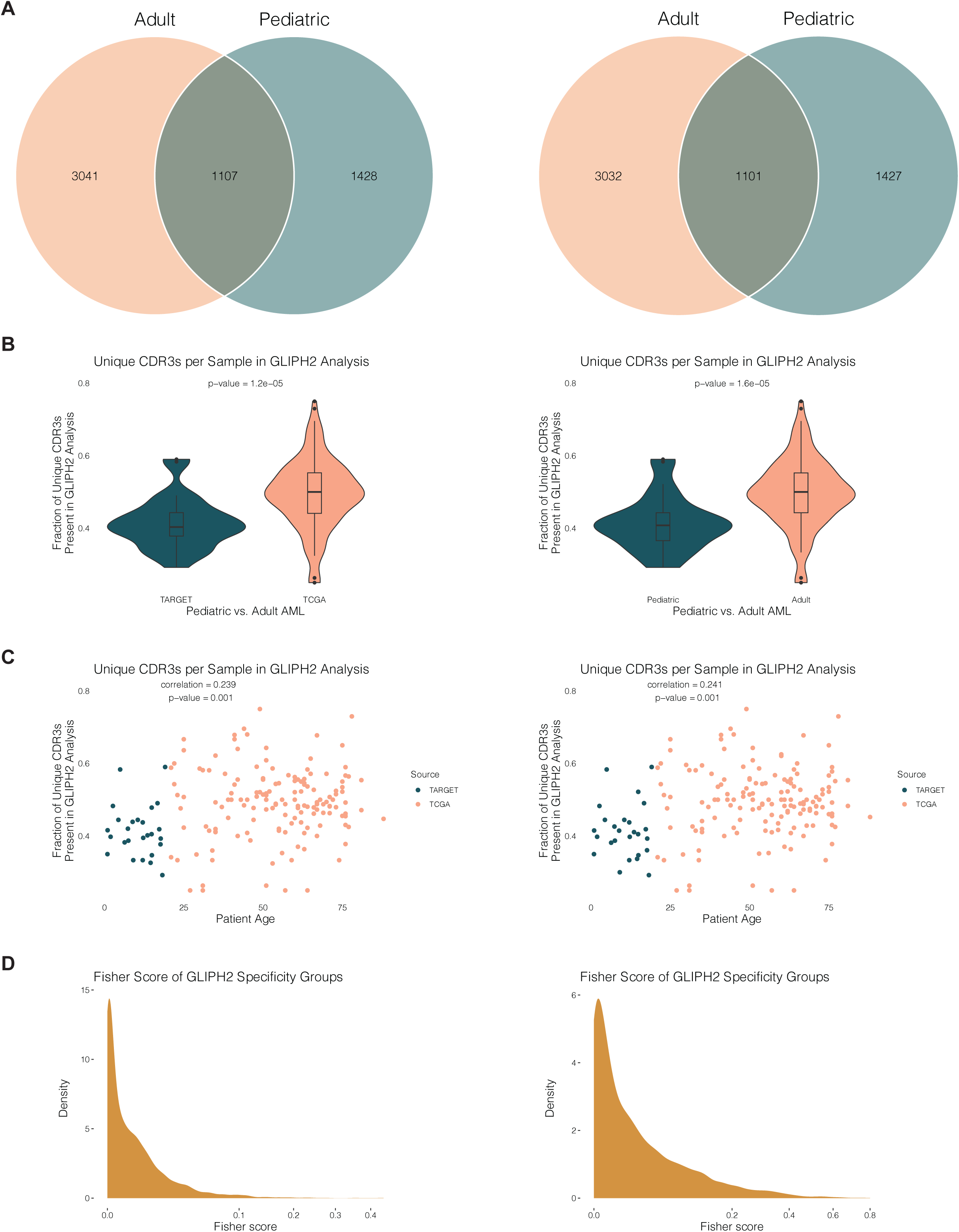
Comparison between adult and pediatric AML. **A)** Venn diagram depicting intersection of TCR specificity groups among adult and pediatric AML patients. We inferred TCR specificities by applying GLIPHII to TCRs derived from RNA-seq datasets (TCGA and TARGET). **B)** Violin plots showing fraction of unique TCR sequences grouped in TCR specificities by GLIPHII in adult vs pediatric AML patients (two-sided Student’s t-test) **C)** Scatter plot showing associations between fraction of TCR sequences grouped by GLIPHII as a measure of TCR convergence, and age at diagnosis across pediatric and adult AML patients. **D)** Histogram showing distribution of Fisher scores derived from GLIPHII inference analyses. In all panels, TCR specificities were inferred using GLIPHII default (left panels) as well as a custom pediatric reference dataset (right panels).

**Figure S6.**
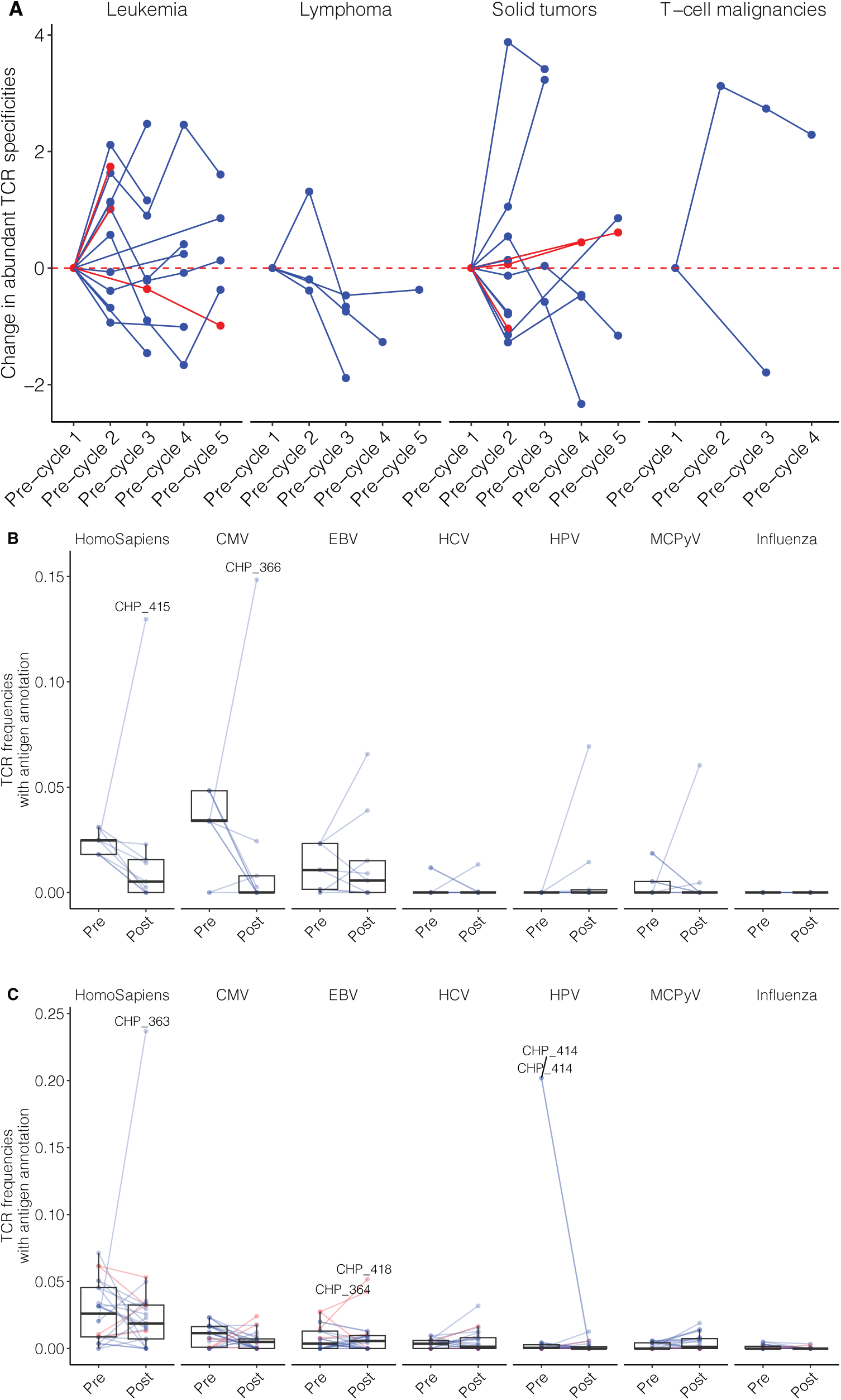
**A)** Spider plots depicting patient-specific changes in Simpson diversity of TCR specificity groups inferred by GLIPHII over time relative to pre-treatment samples in leukemia, lymphoma, and solid tumors plots. Each line corresponds to one patient and y-axes show scaled change relative to pre-therapy samples. Red color denotes relapsed disease, while blue color show patients with complete remission. **B)** Boxplot showing frequency of TCRs within specificity groups that were annotated with human or viral antigens across paired pre- and post-therapy lymphoma samples. **C)** Boxplot showing frequency of TCRs within specificity groups that were annotated with human or viral antigens across paired pre- and post-therapy solid tumor samples. In boxplots, boxes show median and interquartile range (IQR) and whiskers represent 1.5 × IQR.

**Figure S7.**
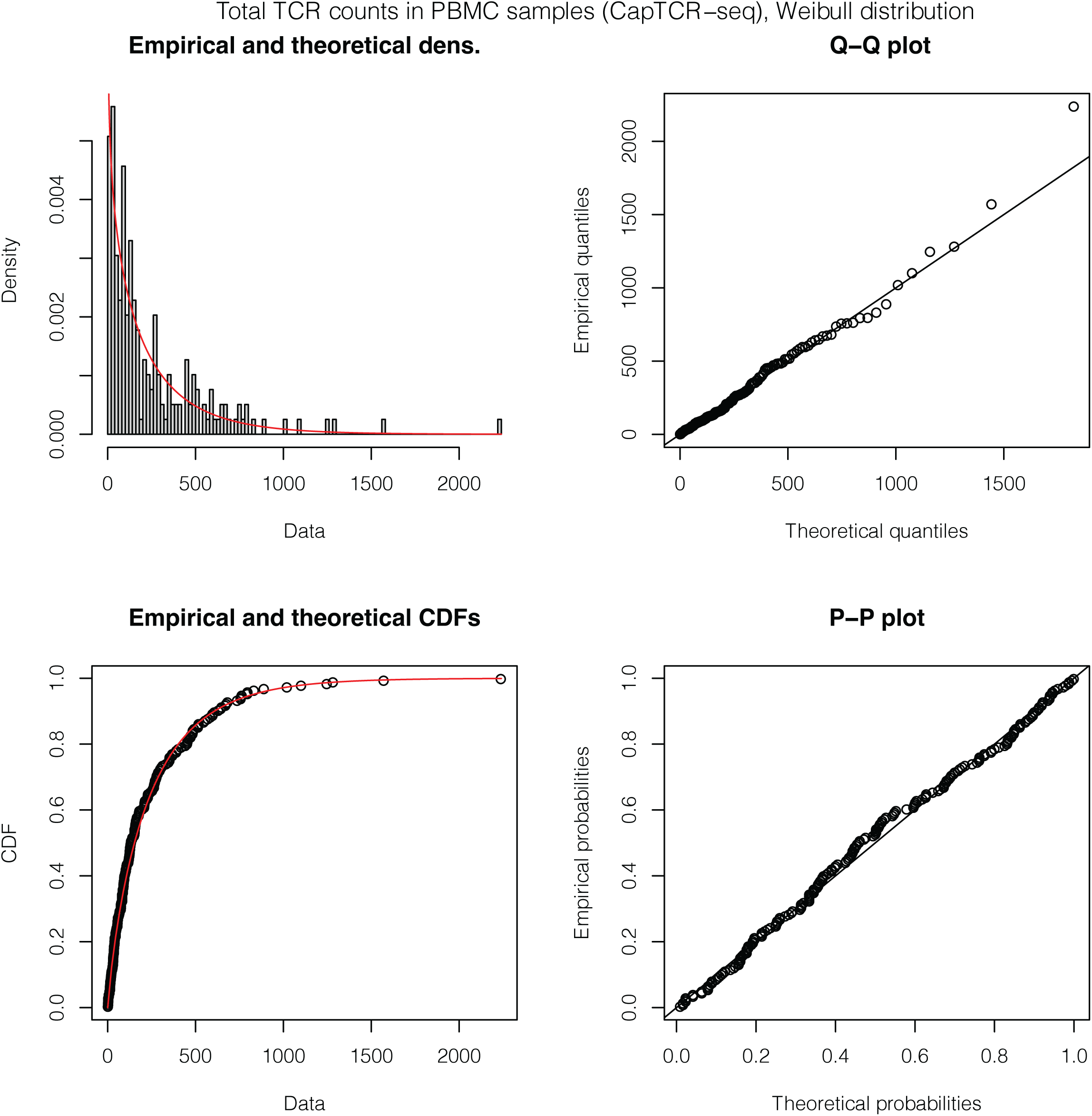
Distribution of TCRs profiled using CapTCR-seq. **A)** Histogram showing distribution of TCR frequencies. The red line shows the fitted Weibull distribution. **B-D)** Goodness-of-fit plots for the fitted Weibull distribution, which was used for random sampling of healthy controls. (Q-Q plot (B), Cumulative Distribution Function (CDF) plot for empirical data and the fitted model (C), and P-P plot (D).

## Supplementary Tables

**TableS1.** Details of Patient and Sample Characteristics and Their Immune Profiling Data.

**TableS2.** Univariate Logistic Regression Analysis of Immune Features and Relapse at Pre-Therapy.

**TableS3.** Univariate Logistic Regression Analysis of Immune Feature Changes During Therapy and Relapse.

**TableS4.** Logistic Regression Analysis of DIS and Relapse.

**TableS5.** ERGO-II Predictions of TCR-Peptide Binding for Public TCR Sequences.

**TableS6.** GLIPHII Results for the Pediatric Cancer Samples and TCR Sequences with Known Specificities.

**TableS7.** TCR Specificity Network Analysis Results for 22 Components Present in Pre- and Post-Therapy Samples.

## References

1. Alexander, S. et al. Prevention of Bacterial Infection in Pediatric Oncology: What Do We Know, What Can We Learn? Pediatr Blood Cancer 59, 16–20 (2012).

2. Vento, S. & Cainelli, F. Infections in patients with cancer undergoing chemotherapy: aetiology, prevention, and treatment. The Lancet Oncology 4, 595–604 (2003).

3. Whittle, S. B., Williamson, K. C. & Russell, H. V. Incidence and risk factors of bacterial and fungal infection during induction chemotherapy for high-risk neuroblastoma. Pediatric Hematology and Oncology 34, 331–342 (2017).

4. Guilcher, G. M. T. et al. Immune function in childhood cancer survivors: a Children’s Oncology Group review. The Lancet Child & Adolescent Health 5, 284–294 (2021).

5. Williams, A. P. et al. Immune reconstitution in children following chemotherapy for acute leukemia. eJHaem 1, 142–151 (2020).

6. Mackall, C. L. et al. Lymphocyte Depletion During Treatment With Intensive Chemotherapy for Cancer. Blood 84, 2221–2228 (1994).

7. Mackall, C. L. et al. Distinctions between CD8+ and CD4+ T-cell regenerative pathways result in prolonged T-cell subset imbalance after intensive chemotherapy. Blood 89, 3700–3707 (1997).

8. Mustafa, M. M. et al. Immune Recovery in Children With Malignancy After Cessation of Chemotherapy. Journal of Pediatric Hematology/Oncology 20, 451 (1998).

9. Yao, Z. et al. Recovery of lymphocyte subpopulations is incomplete in the long-term setting in pediatric solid tumor survivors. Pediatrics International 64, e15257 (2022).

10. van Tilburg, C. M. et al. Immune reconstitution in children following chemotherapy for haematological malignancies: a long-term follow-up. British Journal of Haematology 152, 201–210 (2011).

11. Rabin, K. R. et al. Absolute lymphocyte counts refine minimal residual disease-based risk stratification in childhood acute lymphoblastic leukemia. Pediatric Blood & Cancer 59, 468–474 (2012).

12. Galvez-Silva, J. et al. Prognostic Analysis of Absolute Lymphocyte and Monocyte Counts after Autologous Stem Cell Transplantation in Children, Adolescents, and Young Adults with Refractory or Relapsed Hodgkin Lymphoma. Biology of Blood and Marrow Transplantation 23, 1276–1281 (2017).

13. Geoerger, B. et al. Atezolizumab for children and young adults with previously treated solid tumours, non-Hodgkin lymphoma, and Hodgkin lymphoma (iMATRIX): a multicentre phase 1–2 study. The Lancet Oncology 21, 134–144 (2020).

14. Geoerger, B. et al. Pembrolizumab in paediatric patients with advanced melanoma or a PD-L1-positive, advanced, relapsed, or refractory solid tumour or lymphoma (KEYNOTE-051): interim analysis of an open-label, single-arm, phase 1–2 trial. The Lancet Oncology 21, 121–133 (2020).

15. Davis, K. L. et al. Nivolumab in children and young adults with relapsed or refractory solid tumours or lymphoma (ADVL1412): a multicentre, open-label, single-arm, phase 1–2 trial. The Lancet Oncology 21, 541–550 (2020).

16. Chen, G. M. et al. Integrative Bulk and Single-Cell Profiling of Premanufacture T-cell Populations Reveals Factors Mediating Long-Term Persistence of CAR T-cell Therapy. Cancer Discov 11, 2186–2199 (2021).

17. Das, R. K., Vernau, L., Grupp, S. A. & Barrett, D. M. Naïve T-cell Deficits at Diagnosis and after Chemotherapy Impair Cell Therapy Potential in Pediatric Cancers. Cancer Discov 9, 492–499 (2019).

18. Singh, N., Perazzelli, J., Grupp, S. A. & Barrett, D. M. Early memory phenotypes drive T cell proliferation in patients with pediatric malignancies. Science Translational Medicine 8, 320ra3–320ra3 (2016).

19. McNeer, J. L., Rau, R. E., Gupta, S., Maude, S. L. & O’Brien, M. M. Cutting to the Front of the Line: Immunotherapy for Childhood Acute Lymphoblastic Leukemia. Am Soc Clin Oncol Educ Book e132–e143 (2020) doi:10.1200/EDBK_278171.

20. Diorio, C. & Maude, S. L. CAR T cells vs allogeneic HSCT for poor-risk ALL. Hematology 2020, 501–507 (2020).

21. Mulder, D. T. et al. CapTCR-seq: hybrid capture for T-cell receptor repertoire profiling. Blood Adv 2, 3506–3514 (2018).

22. Parsons, E. et al. Regulatory T Cells in Endemic Burkitt Lymphoma Patients Are Associated with Poor Outcomes: A Prospective, Longitudinal Study. PLoS One 11, e0167841 (2016).

23. Pauken, K. E. et al. The PD-1 Pathway Regulates Development and Function of Memory CD8+ T Cells following Respiratory Viral Infection. Cell Rep 31, 107827 (2020).

24. Gerlach, C. et al. Heterogeneous differentiation patterns of individual CD8+ T cells. Science 340, 635–639 (2013).

25. Freeman, G. J. et al. Engagement of the PD-1 immunoinhibitory receptor by a novel B7 family member leads to negative regulation of lymphocyte activation. J Exp Med 192, 1027–1034 (2000).

26. Kenswil, K. J. G. et al. Immune composition and its association with hematologic recovery after chemotherapeutic injury in acute myeloid leukemia. Experimental Hematology 105, 32–38.e2 (2022).

27. Josse, J. & Husson, F. missMDA: A Package for Handling Missing Values in Multivariate Data Analysis. Journal of Statistical Software 70, 1–31 (2016).

28. Springer, I., Tickotsky, N. & Louzoun, Y. Contribution of T Cell Receptor Alpha and Beta CDR3, MHC Typing, V and J Genes to Peptide Binding Prediction. Front Immunol 12, 664514 (2021).

29. Huang, H., Wang, C., Rubelt, F., Scriba, T. J. & Davis, M. M. Analyzing the Mycobacterium tuberculosis immune response by T-cell receptor clustering with GLIPH2 and genome-wide antigen screening. Nature Biotechnology 38, 1194–1202 (2020).

30. Zhang, J. et al. Immune receptor repertoires in pediatric and adult acute myeloid leukemia. Genome Medicine 11, 73 (2019).

31. Mitchell, A. M. et al. Temporal development of T cell receptor repertoires during childhood in health and disease. JCI Insight 7, (2022).

32. Shugay, M. et al. VDJdb: a curated database of T-cell receptor sequences with known antigen specificity. Nucleic Acids Research 46, D419–D427 (2018).

33. Soleimani S, et al. T-cell receptor repertoire in cell-free DNA as a proxy for tumour infiltrates in patients treated with pembrolizumab. In revision.

34. Emerson, R. O. et al. Immunosequencing identifies signatures of cytomegalovirus exposure history and HLA-mediated effects on the T cell repertoire. Nat Genet 49, 659–665 (2017).

35. Huuhtanen, J. et al. Evolution and modulation of antigen-specific T cell responses in melanoma patients. Nat Commun 13, 5988 (2022).

36. Erickson, J. J. et al. Viral acute lower respiratory infections impair CD8^+^ T cells through PD-1. J Clin Invest 122, 2967–2982 (2012).

37. Li, Y. et al. Impact of T-cell immunity on chemotherapy response in childhood acute lymphoblastic leukemia. Blood 140, 1507–1521 (2022).

38. Kanakry, C. G. et al. Early lymphocyte recovery after intensive timed sequential chemotherapy for acute myelogenous leukemia: peripheral oligoclonal expansion of regulatory T cells. Blood 117, 608–617 (2011).

39. Knaus, H. A. et al. Signatures of CD8+ T cell dysfunction in AML patients and their reversibility with response to chemotherapy. JCI Insight 3, e120974.

40. Chiou, S.-H. et al. Global analysis of shared T cell specificities in human non-small cell lung cancer enables HLA inference and antigen discovery. Immunity 54, 586–602.e8 (2021).

41. Kroemer, G., Galluzzi, L., Kepp, O. & Zitvogel, L. Immunogenic Cell Death in Cancer Therapy. Annual Review of Immunology 31, 51–72 (2013).

42. Krysko, D. V. et al. Immunogenic cell death and DAMPs in cancer therapy. Nat Rev Cancer 12, 860–875 (2012).

43. Wong, D. et al. Early Cancer Detection in Li–Fraumeni Syndrome with Cell-Free DNA. Cancer Discovery 14, 104–119 (2024).

44. Bratman, S. V. et al. Personalized circulating tumor DNA analysis as a predictive biomarker in solid tumor patients treated with pembrolizumab. Nat Cancer 1, 873–881 (2020).

45. Wang, T. T. et al. High efficiency error suppression for accurate detection of low-frequency variants. Nucleic Acids Research 47, e87–e87 (2019).

46. Bolotin, D. A. et al. MiXCR: software for comprehensive adaptive immunity profiling. Nat. Methods 12, 380–381 (2015).

47. Sun, X. et al. Longitudinal analysis reveals age-related changes in the T cell receptor repertoire of human T cell subsets. J Clin Invest 132, (2022).

48. Chao, A. et al. Rarefaction and extrapolation with Hill numbers: a framework for sampling and estimation in species diversity studies. Ecological Monographs 84, 45–67 (2014).

49. Yue, T. et al. TCRosetta: an integrated analysis and annotation platform for T-cell receptor sequences. Genomics, Proteomics & Bioinformatics qzae013 (2024) doi:10.1093/gpbjnl/qzae013.

50. Traag, V. A., Waltman, L. & van Eck, N. J. From Louvain to Leiden: guaranteeing well-connected communities. Sci Rep 9, 5233 (2019).

51. Gee, M. H. et al. Antigen Identification for Orphan T Cell Receptors Expressed on Tumor-Infiltrating Lymphocytes. Cell 172, 549–563.e16 (2018).

